# A robust benchmark for detecting low-frequency variants in the HG002 Genome In A Bottle NIST reference material

**DOI:** 10.1101/2024.12.02.625685

**Authors:** Camille A. Daniels, Adetola Abdulkadir, Megan H. Cleveland, Jennifer H. McDaniel, David Jáspez, Luis Alberto Rubio-Rodríguez, Adrián Muñoz-Barrera, José Miguel Lorenzo-Salazar, Carlos Flores, Byunggil Yoo, Sayed Mohammad Ebrahim Sahraeian, Yina Wang, Massimiliano Rossi, Arun Visvanath, Lisa Murray, Wei-Ting Chen, Severine Catreux, James Han, Rami Mehio, Gavin Parnaby, Andrew Carroll, Pi-Chuan Chang, Kishwar Shafin, Daniel Cook, Alexey Kolesnikov, Lucas Brambrink, Mohammed Faizal Eeman Mootor, Yash Patel, Takafumi N. Yamaguchi, Paul C. Boutros, Karolina Sienkiewicz, Jonathan Foox, Christopher E. Mason, Bryan R. Lajoie, Carlos A. Ruiz-Perez, Semyon Kruglyak, Justin M. Zook, Nathan D. Olson

## Abstract

Somatic mosaicism is an important cause of disease, but mosaic and somatic variants are often challenging to detect because they exist in only a fraction of cells. To address the need for benchmarking subclonal variants in normal cell populations, we developed a benchmark containing mosaic variants in the Genome in a Bottle Consortium (GIAB) HG002 reference material DNA from a large batch of a normal lymphoblastoid cell line. First, we used a somatic variant caller with high coverage (300x) Illumina whole genome sequencing data from the Ashkenazi Jewish trio to detect variants in HG002 not detected in at least 5% of cells from the combined parental data. These candidate mosaic variants were subsequently evaluated using >100x BGI, Element, and PacBio HiFi data. High confidence candidate SNVs with variant allele fractions above 5% were included in the HG002 draft mosaic variant benchmark, with 13/85 occurring in medically relevant gene regions. We also delineated a 2.45 Gbp subset of the previously defined germline autosomal benchmark regions for HG002 in which no additional mosaic variants >2% exist, enabling robust assessment of false positives. The variant allele fraction of some mosaic variants is different between batches of cells, so using data from the homogeneous batch of reference material DNA is critical for benchmarking these variants. External validation of this mosaic benchmark showed it can be used to reliably identify both false negatives and false positives for a variety of technologies and detection algorithms, demonstrating its utility for optimization and validation. By adding our characterization of mosaic variants in this widely-used cell line, we support extensive benchmarking efforts using it in simulation, spike-in, and mixture studies.

## Introduction

Germline variant calling in human genome studies typically targets heterozygous and homozygous variants occurring at variant allele fractions (VAFs) of 50% or 100%, respectively. However, variants can occur at lower fractions if they are only present in a subset of cells due to somatic mosaicism, making them harder to detect and requiring different variant calling methods to identify and characterize them. Somatic mutations occur within a genome after conception, are typically not inherited, and are only present in a subset of cells. While many of these mutations are non-pathogenic, others can cause unrestricted cell growth and lead to cancer or play roles in the development of neurodegenerative, monogenic, and complex diseases (Freed, Stevens, and Pevsner 2014; Truty et al. 2023). An initiative by the National Institutes of Health (NIH) Common Fund called Somatic Mosaicism across Human Tissues (SMaHT, https://smaht.org/) has been instituted to establish a repository of mosaic variants from various healthy tissue types and address the lack of resources to study somatic mosaicism. This effort has recognized a need for benchmarks to evaluate and validate low frequency variant calling methods. In this work, we use the term ‘mosaic variants’ to mean variants present in some cells but not all cells in a large batch of DNA from the deeply-characterized Genome in a Bottle (GIAB) normal lymphoblastoid cell line HG002.

Previous GIAB benchmarks using whole-genome sequencing (WGS) have focused on characterizing germline small (Zook et al. 2014, 2016, 2019; Chin et al. 2020; Wagner, Olson, Harris, Khan, et al. 2022; Wagner, Olson, Harris, McDaniel, et al. 2022) and structural (Parikh et al. 2016; Zook et al. 2020; Wagner, Olson, Harris, McDaniel, et al. 2022) variants which generally ignore variants with <30% VAF. These benchmark sets include high confidence variant calls and regions. The benchmark variants are confident homozygous and heterozygous variants in a sample relative to a reference genome (VCF file), and the benchmark regions (BED file) are genomic regions that are confidently identified as homozygous reference or a benchmark variant. The benchmark variants enable users to identify true positives and false negatives in their query callset, and benchmark regions enable the identification of false positive variants.

The reliable identification of errors (RIDE) principle is used to determine if a GIAB benchmark set is fit for purpose, specifically the identifying false positives and false negatives across a variety of high-quality methods (Olson et al. 2023). In addition to the GIAB benchmark sets, the GIAB Consortium has worked with the Global Alliance for Genomics and Health (GA4GH) to define best practices for benchmarking small variants (Krusche et al. 2019). While the GIAB benchmark sets and benchmarking methods have been used to evaluate small and structural variant calling methods, as well as training machine learning and deep learning based variant calling methods, low frequency or mosaic variants in GIAB reference materials have not been previously characterized.

Characterization of mosaic variants in GIAB reference materials would allow researchers to use the GIAB reference materials and genome sequencing data generated from the material to validate mosaic and somatic variant calling methods, as well as other uses such as negative controls when evaluating methods for detecting off-target genome edits. To ensure that NGS protocols and bioinformatic pipelines can accurately and reliably detect low frequency mutations, well-characterized reference samples are needed. Previous efforts developing benchmarks for low frequency variants have used a variety of strategies. For example, data from four historical cell lines were mixed in a variety of ratios to mimic mosaic variants at different fractions (Ha et al. 2023, 2022). However, because these are germline variants, some mosaic and somatic variant callers will filter them, and there is a lack of clarity in accurately identifying true negatives. Therefore, cancer-focused benchmarks have taken other approaches, like injecting synthetic somatic variants into real data (Ewing et al. 2015), simulating tumor subclonality (Salcedo et al. 2024), creating synthetic DNA with somatic mutations spiked into a normal background sample (Sims et al. 2016), engineering normal samples to contain somatic variants (Pfeifer et al. 2022), comparing paired tumor and normal cell lines derived from a single individual (Fang et al. 2021; McDaniel et al. 2024), and creating mixtures of cell lines from different individuals (Jones et al. 2021).

Here, we complement these previous efforts by leveraging publicly available homogenous batches of DNA reference materials linked to explicit consent for public genome data sharing. By characterizing the baseline mosaic variants in this cell line, it can be used more robustly as a negative control or background in many of the other benchmarking approaches (e.g., when modifying reads to contain mutations, adding spike-in DNA with mutations, or mixing with other samples). Specifically, we present an initial mosaic benchmark for the GIAB HG002 reference material from a broadly-consented individual from the Personal Genome Project (Ball et al. 2012). The GIAB reference material (RM) used for this study originates from a large homogenous batch of DNA isolated from the HG002 cell line (Zook et al. 2016). Mosaic variants in this RM DNA may be from somatic mosaicism in the individual’s B cells or from mutations that have arisen during the cell line generation and culturing process. To generate this new benchmark we used a trio-based approach (**Figure 1**). We first identified potential mosaic variants using the 300x coverage Illumina sequencing data and the Strelka2 tumor/normal somatic variant caller with son (HG002) as the tumor sample and parents (HG003 + HG004) as the normal sample. High-coverage orthogonal sequencing data for the NIST HG002 RM DNA was used to validate the low frequency variant calls identified by Strelka2. While there is some preliminary evidence of possible mosaics occurring above 30% VAF in human genome data, this study focused on somatic mosaic variants (≤ 30 % VAF).

**Figure 1.**
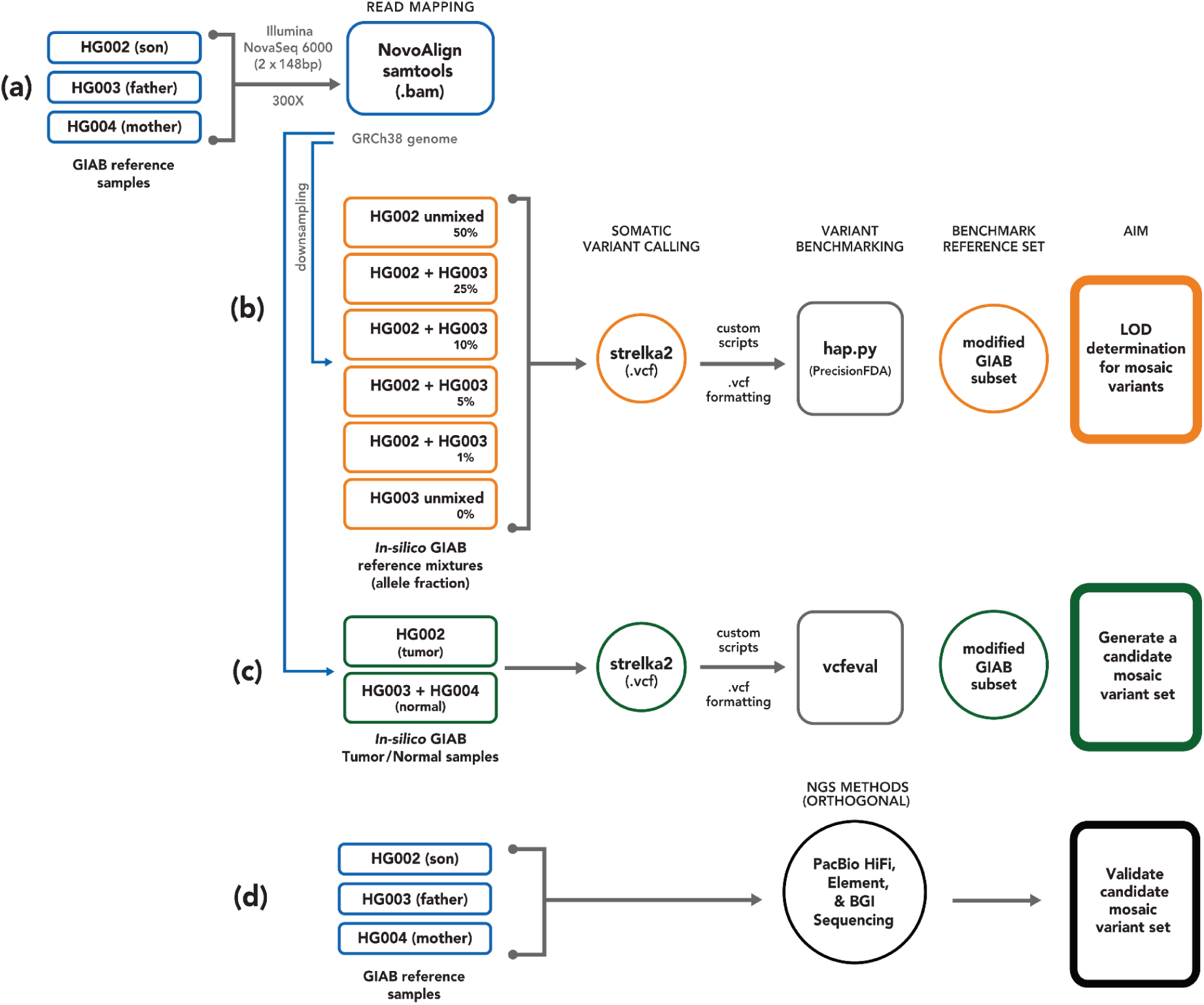
Trio-based methodology using high coverage Illumina data, Strelka2 somatic caller, and orthogonal next generation sequencing datasets for candidate mosaic variant detection and validation in HG002. (A) AJ trio (NIST RM - HG002, HG003, and HG004) sequencing and reference mapping (GRCh38) were initially performed by Zook et al 2016. (B) *In silico* sample mixtures were created using HG002 and HG003, treating HG003 as normal and the mixtures as tumor, to determine the limit of detection for variant allele fraction. Strelka2 somatic calling and benchmarking with hap.py was conducted using the GIAB mixtures to estimate a limit of detection (LOD). (C) To identify potential mosaic and de novo variants, a tumor-normal Strelka2 somatic run, with HG002 (son) as tumor and HG003+HG004 (combined parents) as normal, was performed. (D) The Strelka2 callset was benchmarked against the GIAB v4.2.1 small variant benchmark with vcfeval to create a candidate variant set, and three orthogonal high-coverage short- and long-read sequencing technologies were used for validation.

## Results

### Mosaic benchmark set generation and characterization

The HG002 mosaic benchmark set includes 85 validated and manually curated SNVs and benchmark regions covering 2.45 Gbp (**Figure 2**). To arrive at this benchmark, first, potential mosaic variants in the HG002 NIST RM DNA were identified using the 300× Illumina AJ trio dataset and the Strelka2 somatic variant caller, with HG002 (son) data as tumor and HG003 + HG004 (parents) as normal. Strelka2 callset contained ≈1.27 million passing and filtered SNVs and indels, with 425,679 potential mosaic variants after excluding variants in the AJ Trio v4.2.1 GIAB small variant benchmark. Exclusion of AJ trio v4.2.1 complex and structural variants in HG002 further reduced the number of potential somatic variants to 366,728 (**Supplemental Figure 2**). While only 1,916 SNVs and 21 indels passed the Strelka2 filter, we kept filtered variants for downstream analysis (**Supplemental Table 2**) to reduce the probability of missing variants based on our *in silico* mixture experiments (**Supplemental Figure 1**).

**Figure 2.**
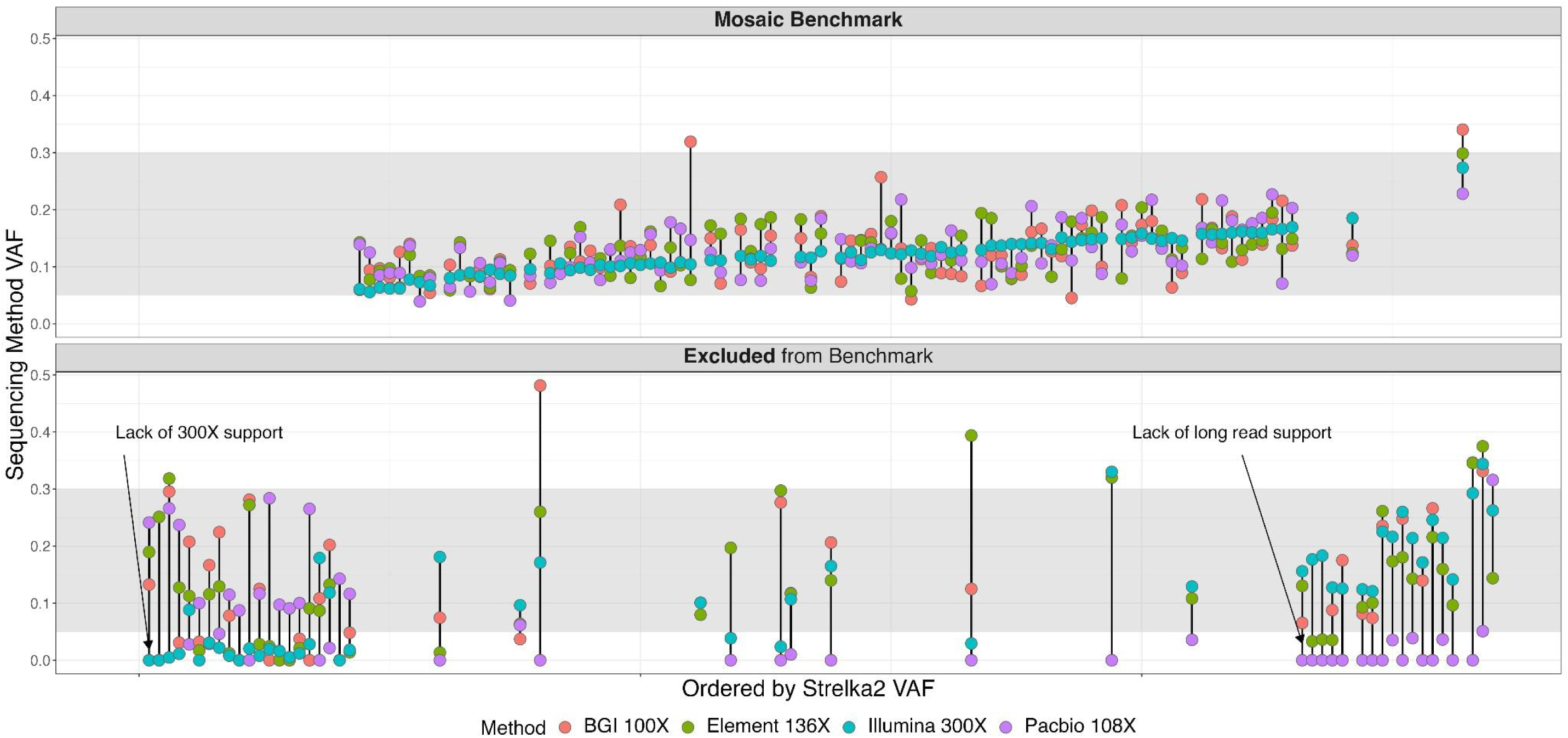
Manually curated potential mosaic variants (135) depicted as vertical lines and arranged by increasing Strelka2 variant allele frequency (X-axis, left to right). Colored dots represent HG002 Illumina 300x (teal) and orthogonal tech datasets for each variant (BGI 100x - red, Element 136x - green, and PacBio HiFi 108x - purple) with corresponding bam-readcount VAFs located on the X-axis. Shaded area indicates the range of VAF (5% to 30%) of variants targeted for inclusion in the benchmark. The top facet illustrates 85 high-confidence SNVs **included** in the HG002 mosaic benchmark v1.0, while the bottom facet shows 50 SNVs **excluded** from the benchmark.

To define our mosaic benchmark variants and regions, we evaluated support for the 366,728 potential mosaic variants across multiple short- and long-read technologies with at least 100x coverage per technology. We created a database with these potential mosaics that included Strelka2 VCF annotations, read support metrics from multiple sequencing technologies, and genomic context (**Data Availability**). Using an initial set of heuristics (**Supplemental Figure 3**), we filtered this database and identified 135 variants for manual curation. After manual curation, an additional 50 variants were excluded due to either lack of long-read support, high coverage short-read support, or low VAF reported by Strelka2 (**Figure 2**). Many of these excluded SNVs were detected in one or both parents (HG003 and/or HG004) and were found in GRCh38 segmental duplication regions associated with mapping errors and copy number variation. To create the HG002 mosaic benchmark v1.0 BED file, we excluded genomic regions with tandem repeats and homopolymers, regions containing variants that could not be confidently determined to be >5% or <2% VAF (**Supplemental Figure 3 - yellow squares**), and regions <50bp, since small benchmark regions can cause problems with benchmarking complex variants.

### Benchmark variant and region characteristics

The HG002 mosaic benchmark v1.0 contains 85 SNVs with VAFs between 5% to 30% (**Figure 3a**, 99% one-sided confidence interval ≥ 0.05, calculated based on the combined read support across multiple orthogonal sequencing technologies, **Figure 2, Supplemental Figure 5, Data Availability**). The mosaic benchmark included 2.45 Gbp (89.5% of the GRCh38 non-gapped assembled bases in the autosomes) with 891,240 regions, ranging from 50 bp - 76.6 kbp (**Data Availability**). The benchmark set included variants in challenging genomic regions with two of the 85 variants in homopolymers and two in low mappability regions (**Figure 3b**). While no mosaic benchmark variants were observed in GRCh38 coding regions, 13 of 85 (∼15%) variants are in medically relevant genes (Wagner, Olson, Harris, McDaniel, et al. 2022), and benchmark regions included > 90% of the bases in these genes (**Supplemental Table 4**). The HG002 mosaic benchmark regions include a total of 417,752,807 bases in medically relevant genes, and included > 90% of bases in 3,871 medically relevant genes.

**Figure 3.**
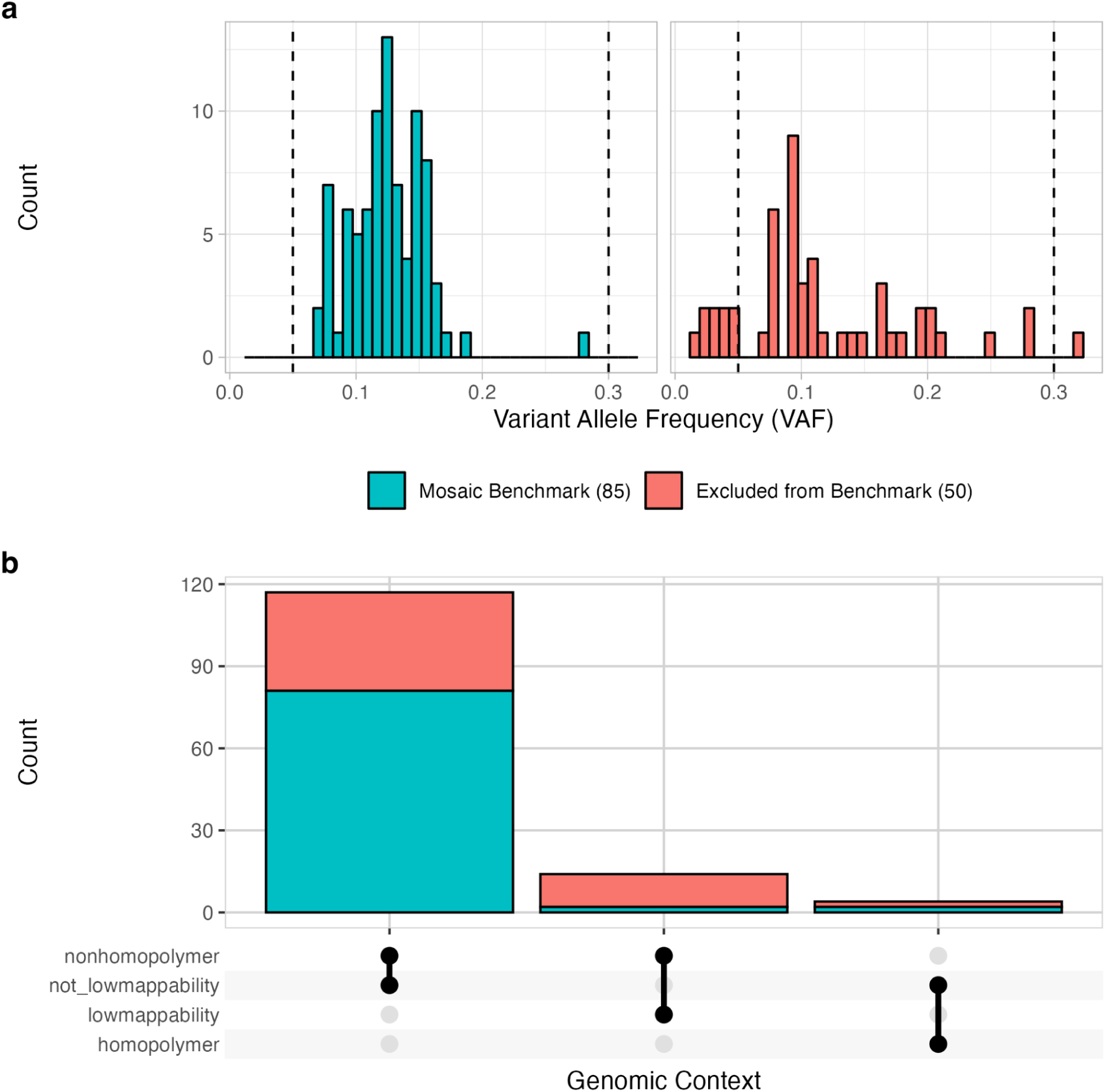
SNV variant allele fractions (VAF) (**a**) for HG002-GRCh38 mosaic benchmark v1.0 and manually curated variants excluded from the benchmark. Values represent VAFs combined across all orthogonal technologies (BGI, Element, Illumina, PacBio Revio, and Sequel). Dashed vertical lines represent the targeted VAF range (5% - 30%) for the HG002 mosaic benchmark. Manually curated variant counts based on GIAB GRCh38 genome stratifications (**b**) reveal most mosaic benchmark v1.0 variants occur in easy-to-map and non-homopolymer regions of the genome.

### Mosaic Variants Reveal GIAB Material Batch Effects

We identified differing VAF profiles for the benchmark mosaic variants between the large batch of DNA distributed as NIST RM 8391 and various unknown batches of non-reference material DNA from Coriell (GM24385/NA24385, **Supplemental Table 1**). For both Element and PacBio Revio, the mosaic benchmark VAFs were significantly higher than datasets generated using non-RM DNA, so that these differences are likely to result from changes in the cell line rather than random sampling (**Figure 4**). The larger difference for PacBio Revio suggests additional clonal changes prior to its sequencing. While variants differed in VAF between batches, they generally were present in all batches.

**Figure 4.**
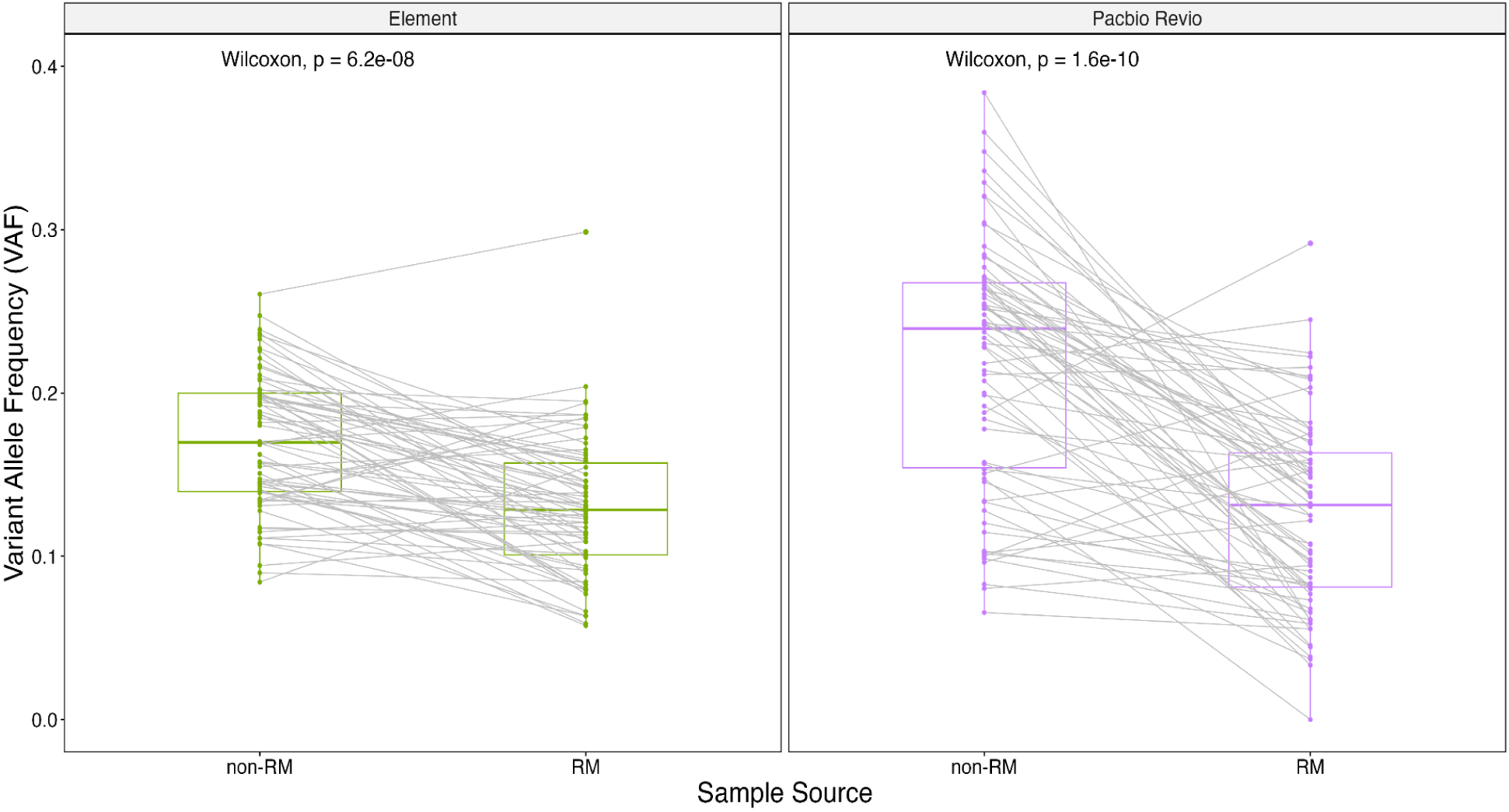
Mosaic variants change VAF between batches of DNA. HG002 mosaic benchmark variant allele fractions (VAFs) for NIST reference material (RM) 8391 and different batches of non-RM DNA (Coriell, NA24385) for two orthogonal technologies (Element and PacBio Revio). Higher VAFs were observed in direct VAF comparisons between materials compared to GIAB reference material. Coverage: Element RM: 136x, non-RM: 100x; PacBio Revio RM: 48x, non-RM: 120x.

#### External Validation

Eight somatic variant calling groups submitted callsets for use in validating the draft mosaic benchmark. Most groups used the same GIAB HG002 Illumina 300x data used to generate the mosaic v1.0 benchmark, one group provided an in-house HG002 40x callset, and others produced additional short-(Element 70x and 100x, PacBio Onso 35x) and long-read (PacBio Revio 130x) HG002 callsets (**Supplemental Table 5**). The evaluations focused on curating differences between external callsets and a draft version of the mosaic benchmark generated from both HG002 RM (NIST RM 8391) and non-RM (Coriell, NA24385) data. Unlike the final benchmark, the draft benchmark included some non-RM data because RM data from some technologies were not yet available.

During the evaluation, we found that the draft mosaic benchmark variants were reliable, but some variants were incorrectly filtered by our initial heuristics. For example, we had initially ignored candidate mosaic variants with lower than normal coverage, because low coverage regions generally were associated with alignment errors around larger germline variants. However, some true mosaic variants were ignored but kept in the benchmark regions, so we modified the heuristics for v1.0 to only ignore variants below the 0.5% quantile in coverage for all datasets combined or HiFi only. Additionally, the draft benchmark had missed some variants substantially different in VAF between RM and non-RM DNA, so we only used RM DNA data for the v1.0 benchmark.

We observed a single case of a true mosaic SNV missed by our trio-based approach, identified by one method using only HG002 data (chr10:106867519). Interestingly, this variant appears to be a heterozygous germline variant in the father (HG003), but HG002 inherited the reference allele from HG003 and HG004, so it appears to be a true mosaic variant in HG002 that matches the father’s variant but occurred independently (**Supplemental Figure 5**), which has been seen previously (Fasching et al. 2021). This variant and the adjacent 50bps on either side were not included in the benchmark VCF or BED files, so the parents can be used as normal when evaluating tumor/normal somatic variant callers. It is possible there may be additional mosaic variants missed by the benchmark like this if they coincide with parental germline variants, but this was the only one identified during the external evaluation.

The external evaluation results informed refinement of the draft benchmark into v1.0, which considered only NIST RM 8391 data adding 15 mosaic SNVs no longer filtered by our heuristics, for a total of 85 HG002 mosaic benchmark v1.0 variants. Using hap.py to compare the external callsets to the v1.0 benchmark, a majority of mosaic benchmark variants (≥ 87%) were present in short- and long-read callsets by the six groups that used high-coverage datasets. Most variants in the external validation callsets not present in the v1.0 benchmark as identified by hap.py had VAFs below 5%, our established limit of detection, suggesting that the HG002 mosaic benchmark v1.0 reliably identifies false positives. We further curated 163 SNVs with VAF between 5 and 30% in external callsets but not in the draft benchmark and found that all but two were likely mapping errors due to segmental duplications or copy number variants in HG002, systematic sequencing errors, local alignment errors around germline insertions, or batch effects (for callsets from non-RM sequencing data)(**Supplemental Tables 6**). One SNV (chr6:150458314) is likely a true mosaic variant near a 4bp insertion on the other haplotype that was missed by our mosaic benchmark, but present in callsets from all eight external validator groups. The other remaining variant (chr1:242208421) is likely a true mosaic variant though likely <5% VAF in the RM DNA. These two variants along with 50bp flanking sequences were removed from the mosaic v1.0 bed to generate HG002 mosaic benchmark v1.1.

## Discussion and Conclusions

GIAB benchmark sets have focused on germline variants, which occur 50% or 100% VAF, higher than typical mosaic or somatic variants. To address the need for a benchmark for variants with lower VAFs, which occur in only a fraction of the cells, we developed the first GIAB mosaic SNV benchmark for the highly characterized HG002 genome using data from the GIAB AJ trio (son and parents). We substituted a combined parental BAM for the typical matched normal sample (**Figure 1c**), and followed best practices for somatic variant calling (Koboldt 2020) and developing benchmarks (Olson et al. 2023) by filtering out normal variants and artifacts, performing manual curation to confirm the candidate mosaic set, and comparing the benchmark with orthogonal methods.

We identified 85 benchmark SNVs in the HG002 reference material DNA with VAFs between 5% and 30% by high-coverage Illumina and orthogonal short- and long-read methods (BGI, Element, and PacBio HiFi) (**Figure 2**). External validation confirmed that the benchmark set can be used to reliably identify errors in somatic and mosaic variant callsets. Variants were included in the HG002 mosaic benchmark v1.0 for several reasons: a) passed decision tree heuristics that required sufficient confidence the variants were >5% VAF to be included or <2% VAF to be ignored, b) support in long reads for difficult-to-map regions, c) confirmation by manual curation, and d) typical coverage of the region.

While variants <5% VAF were not retained in this mosaic benchmark due to the limit of detection (LOD) of our discovery approach, further investigation using library preparation that corrects DNA damage due to extraction protocols (Chen et al. 2017), deeper sequencing, high-accuracy sequencing technologies, unique molecular identifiers, and/or incorporation of low fraction somatic variant callers (Xiang et al. 2023) are needed to assess if these lower fraction variants should be included in a future HG002 mosaic benchmark version.

External validation of the benchmark confirmed that the benchmark meets GIAB’s RIDE principle across a variety of somatic and mosaic variant callers. Specifically, manual curation determined it reliably identifies both false positives and false negatives across variant calling methods and sequencing technologies.

While DNA batch effects have not previously been reported in GIAB samples when looking at germline variants, we identified substantial batch effects for mosaic variants when comparing HG002 NIST RM 8391 and non-RM (Coriell - NA24385) using mosaic VAF data. Given the higher VAFs found for mosaic benchmark variants in sequencing datasets generated using non-RM DNA, we suggest using datasets generated using NIST HG002 RM DNA to perform mosaic benchmarking.

We envision the HG002 mosaic benchmark as a GIAB somatic resource applied in use cases including, but not limited to: 1) benchmarking mosaic variant callers, 2) as negative controls for either WGS somatic callers or targeted clinical sequencing in tumor-only mode, 3) benchmarking somatic variant callers in tumor-normal mode using GIAB mixtures, 4) as a dataset which germline researchers can use to filter low fraction somatic variants from their data, and 5) benchmarking for some types of off-target genome edits. Currently, the most commonly used small variant benchmarking methods, e.g., hap.py and rtg vcfeval, do not consider VAF or properly handle differences in how low frequency variants are represented in VCFs. Benchmarking methods development and community defined best practices will significantly improve the utility of the mosaic benchmark set and, in turn, low frequency variant calling performance.

The Medical Device Innovation Consortium (MDIC) is a public-private partnership with the aim of advancing regulatory science for the development and assessment of medical devices. MDIC launched the [Somatic Reference Sample (SRS) Initiative](https://mdic.org/wp-content/uploads/2022/10/MDIC-Initiative-SRS_CS_FINAL-Web.pdf) to develop reference samples that can be widely distributed, so that all stakeholders can have access to the same reference samples. To meet this goal, the SRS Initiative intends to genetically engineer the well-characterized GIAB HG002 genome (Zook et al. 2016) with somatic variants. To establish a baseline for the engineered cell lines, the mosaic benchmark set generated in this study using the unedited AJ trio genomes will be used to assess and validate on and off-target edits of clinically relevant cancer variants in HG002.

We report a new HG002 mosaic variant benchmark v1.1 generated using a trio-based framework with GIAB RM and Strelka2 to produce a high-confidence mosaic call set. This benchmark is important given growing interest from the research community in understanding somatic mosaicism, such as the NIH Common Fund project Somatic Mosaicism Across Human Tissues (SMaHT at https://smaht.org/). This HG002 mosaic benchmark will also serve as the genomic background for the upcoming SRS Initiative, a MDIC project, focused on providing RMs to improve cancer and disease diagnostics.

## Methods

### Reference Material High-Coverage Whole Genome Datasets

High coverage (>100x) whole genome sequencing datasets were generated using three different short-read and one long-read sequencing platforms. High-coverage (300x) Illumina short-read data, previously described (Zook et al. 2016), was used to identify potential mosaic variants. Briefly, six vials of NIST reference gDNA from the GIAB Ashkenazi Jewish (AJ) trio cell lines (HG002- son, HG003- father, HG004- mother; **Figure 1a**) were extracted and Illumina TruSeq PCR-free kit was used to generate paired-end libraries. DNA from each replicate library was multiplexed and sequenced with 2×148bp on an Illumina HiSeq2500 v1 in Rapid mode at NIST (Gaithersburg, MD). Pooled reads from each GIAB sample were mapped to GRCh38 genome (GCA_000001405.15, hs38d1) using NovoAlign v3.02.07 and BAM files were generated, sorted, and indexed with SAMtools v0.1.18. High coverage reads (300x) from the GIAB AJ Trio (NBCI Biosample SAMN03283347, SAMN03283345, SAMN03283346; PGP huAA53E0, hu6E4515, hu8E87A9) were retained for subsequent processing in this study (**Table 1**, **Figure 1a)**.

**Table 1.** GIAB AJ trio (HG002 - son, HG003 - father, and HG004 - mother) and orthogonal datasets for HG002 mosaic benchmark generation. Sequencing data was generated using NIST reference material DNA (NIST RM 8391 for HG002 and NIST RM 8392 trio). See supplemental table 6 for non-reference material (Coriell - NA24385) datasets.

| Library | NIST ID | Coverage | SRA<br>Accession # | Publication |
| --- | --- | --- | --- | --- |
| Illumina<br>2x148bp | HG002 | 300x | SRX847862-SRX848005 | Zook et al.<br>2016 |
| Illumina<br>2x148bp | HG003 | 300x | SRX848006-SRX848173 | Zook et al.<br>2016 |
| Illumina<br>2x148bp | HG004 | 300x | SRX848174-SRX848317 | Zook et al.<br>2016 |
| BGI<br>2x150bp | HG002 | 100x | SRX22242218 | This study |
| Element<br>2x150bp,<br>**standard** | HG002 | 81x | SRR30945810 | This study |
| Element<br>2x150bp,<br>**long**<br>insert | HG002 | 55x | SRR30945809 | This study |
| PacBio<br>HiFi<br>**Sequel**<br>10kb | HG002 | 30x | SRX5327410 | Wenger et al.<br>2019 |
| PacBio<br>HiFi<br>**Sequel**<br>15kb | HG002 | 28x | SRX6908796,<br>SRX6908797,<br>SRX6908798 | Unpublished |
| PacBio<br>HiFi<br>**Revio**<br>20kb | HG002 | 48x | NA | Unpublished* |

\* Data publicly available on the PacBio website (https://www.pacb.com/connect/datasets/) as well as GIAB FTP site (https://ftp-trace.ncbi.nlm.nih.gov/ReferenceSamples/giab/data/AshkenazimTrio/HG002_NA24385_son/PacBio_HiFi-Revio_20231031).

Additional high-coverage HG002 orthogonal datasets were used for mosaic variant validation: BGI (100x), Element (standard and long insert - 136X total), and PacBio HiFi (Sequel and Revio-106x total). High coverage PCR-free 2×150bp BGI data was generated by BGI using the DNBSEQ sequencing platform and basecalled using the DNBSEQ basecalling software with default parameters. Reads were aligned to the hg38 genome (https://hgdownload.cse.ucsc.edu/goldenPath/hg38/bigZips/latest/hg38.fa.gz) using bwa-mem v0.7.17. For Element, standard (400bp) and long insert (1.3 kb) libraries were prepared with 1μg gDNA of HG002 NIST RM, the Kapa Hyper Prep Kit, and Kapa UDI. Seven rounds of Covaris shearing were performed on the DNA libraries, which were subsequently treated with USER enzyme and circularized with Adept Rapid PCR Free workflow using the Element Library Compatibility Kit v1. In a separate run for each insert size, 2×150bp libraries were sequenced across both flow cell lanes on the Element Biosciences AVITI™ platform. Reads from each run were mapped to GRCh38-GIABv3 (https://ftp-trace.ncbi.nlm.nih.gov/ReferenceSamples/giab/data/AshkenazimTrio/HG002_NA24385_son/Element_AVITI_20231018/) using bwa-mem v0.7.17 and SAMtools v1.18 was used to assess mapping results for each insert size. All PacBio libraries were created and sequenced at Pacific Biosciences. Two libraries were previously generated (Wenger et al. 2019), which sheared HG002 NIST RM gDNA using a Megaruptor and 20kb protocol. Size selection was performed on a sageELF DNA system (Sage Science) to capture 10kb and 15kb bands for sequencing. Band sizes were verified on an Agilent 2100 BioAnalyzer using a DNA 12000 kit used as input into the SMRTbell® prep kit 3.0, and sequenced to 30x and 28x-fold coverage, respectively, on a PacBio Sequel System. Default parameters were used to generate consensus (CCS) reads. Using pbmm2 with CCS presets, reads were subsequently aligned to GRCh38_no_alt_analysis reference and phased with WhatsHap v0.7. A PacBio Revio library was created (unpublished, 2023) by shearing 6μg HG002 NIST RM gDNA with a Megaruptor 3 and using 4.5μg as input for the SMRTbell® prep kit 3.0. PippinHT size selection followed along with a 10kb cut. PacBio Revio polymerase and sequencing kits were used for sample prep for sequencing to 48x-fold coverage at Pacific Biosciences, and two SMRT cells were subsequently run for the HG002 sample (24 hour movies). Using the PacBio HiFi-human-WGS-WDL pipeline (https://github.com/PacificBiosciences/HiFi-human-WGS-WDL), HiFi reads were aligned independently to GRCh38-GIABv3 (https://ftp-trace.ncbi.nlm.nih.gov/ReferenceSamples/giab/data/AshkenazimTrio/HG002_NA24385_son/PacBio_HiFi-Revio_20231031/).

### In Silico Mixtures to Assess Limit of Detection

A minimum VAF of 5% for variants to include in the mosaic benchmark set was identified using in-silico mixtures of HG002 and HG003. AJ trio Illumina WGS 300x datasets were subset to chromosome 20 and downsampled using SAMtools (v1.9) to create simulated HG002 + HG003 samples with six different allele fractions (AFs), HG002 unmixed (**50% AF**), 50% HG002 + 50% HG003 (**25% AF**), 20% HG002 + 80% HG003 (**10% AF**), 10% HG002 + 90% HG003 (**5% AF**), 2% HG002 + 98% HG003 (**1% AF**), and HG003 unmixed (**0% AF**). Variant calling was performed with the Strelka2 app on DNAnexus® (Strelka v2.8.4 and SAMtools v1.5) using hs37d5 as the reference. Callsets were benchmarked with hap.py in the precisionFDA app and analyzed to ascertain a limit of detection (LOD; **Figure 1b**). Recall was observed for all SNVs at 99% and 97% for passing SNVs down to 5% VAF, and respectively, fell to 25% and 8% approaching 1% VAF (**Supplemental Figure 1**). Based on these preliminary results, we targeted variants with VAFs > 5% and <30% for inclusion in the HG002 mosaic benchmark set. The GIAB v4.2.1 small variant benchmark only includes variants down to 30% VAF.

### Mosaic Benchmark Set Generation

For the HG002 mosaic benchmark set generation, an initial set of mosaic variants was identified using the Strelka2 somatic variant caller and characterizable genomic regions as the intersection of the GIAB HG002, HG003, and HG004 v4.2.1 benchmark regions, excluding complex (i.e., nearby) variants in any sample. Complex variants were excluded as they tend to cause errors due to differences in variant representation. These initial variants and genomic regions were refined based on a set of heuristics defined based on observations from manually curating targeted subsets of the initial mosaic variants.

The initial set of low fraction variants in HG002 (son) were identified with the Strelka2 somatic variant caller in the Strelka2 DNAnexus® app (Strelka v2.9.10 and SAMtools v1.13) and AJ trio 300x WGS data (**Figure 1c**) using the HG002 BAM as tumor, a combined parental BAM (HG003 + HG004) as normal, and the GRCh38 reference including the hs38d1 decoy FASTA, where the job run was split by chromosome. SNV and indel VCFs were concatenated into a single VCF. Variants with a FILTER column value of VARIANT_DETECTED_IN_NORMAL (i.e., HG003 and HG004) were removed using bcftools v1.16, while variants not detected in the normal sample with the Strelka2 tumor allele read (TAR) depth value > 5 were retained. The merged VCF was formatted with custom scripts for downstream analyses.

Next, variants in the GIAB HG002 v4.2.1 small variant benchmark were excluded from the initial set of low fraction variants. The reformatted Strelka2 VCF compared to the v4.2.1 benchmark set using vcfeval v3.11 with the --*squash-ploidy* flag and the intersection of the HG002, HG003, and HG004 v4.2.1 benchmark regions as target regions (**Supplemental Table 1**). The resulting set of low VAF variants were further refined using custom scripts to exclude regions with GIAB AJ trio v4.2.1 complex and structural variants (**Supplemental Figure 2**). From the resulting set of low VAF variants, Strelka2 passing variants are referred to as candidates and non-passing variants as putative mosaic variants.

A database of candidate and potential mosaic variants was generated for the development and application of a set of heuristics used to define the final mosaic benchmark set. The mosaic variants database (**Data Availability**) contained variant information from the Strelka2 VCF, variant support from multiple orthogonal sequencing datasets, and genomic context annotations. Bam-readcount (Khanna et al. 2021) was used to calculate reference and alternate allele read support for HG002 Illumina (300x) and orthogonal HG002-GRCh38 BAMs from three WGS datasets (**Table 1**): BGI (100x), Element (standard and long insert - 136x total), and PacBio HiFi (Sequel and Revio: 106x total). To remove low quality variants, base and mapping quality filters (-b25, -q40) and a minimum threshold of ≥ 2 read support per orthogonal tech for each variant was applied. The read support counts were used to calculate overall VAF estimates and confidence intervals per orthogonal dataset. Binomial confidence intervals (CI) were calculated using binconf R package (Agresti and Coull 1998; Brown, Tony Cai, and DasGupta 2001; Newcombe 2001). Independent orthogonal CIs used a one-sided confidence coefficient of 95%, while the combined orthogonal CI used a one-sided confidence coefficient of 99%. The GIAB GRCh38 genome stratifications (**Supplemental Table 1**) were used to annotate variants in the database with genomic context using bcftools v1.15.

A series of heuristics (**Supplemental Figure 3**) were used to identify variants for manual curation. This process used combined orthogonal CIs, PacBio read depth thresholds, removed variants that overlap germline indels, and partitioned the data by genomic context. In order to be considered for manual curation, we required a ≥ 0.01 (using a 99% one-sided confidence interval) lower CI threshold of the VAF from the combined orthogonal technologies for easy-to-map variants and a ≥ 0.05 (using a 95% one-sided confidence interval) lower PacBio CI threshold for variants in difficult-to-map regions.

The HG002 mosaic SNV benchmark was defined using the following heuristics in a decision tree (**Supplemental Figure 3**). Initial filtering of the mosaic database (366,728) removed variants with a combined orthogonal upper CI ≤ 0.03. Of the passing variants (6,178), those that fell below the 0.5% quantile thresholds for either total number of reads across combined orthogonal methods or PacBio depth were filtered. Next, database variants that overlapped germline indels were removed. Variants with x_ci/n_ci (i.e., total number of variant reads across combined orthogonal methods/total number of reads across combined orthogonal methods) > 0.5 were removed. Database variants that remained were then partitioned into easy-to-map and not easy-to-map (i.e., low mappability) bins. A combined orthogonal lower CI ≥ 0.05 filter was applied to the easy-to-map variants (2,323), with 121 passing and were retained for manual curation. Not easy-to-map variants (1,668) were partitioned into homopolymer and non-homopolymer bins. After applying a PacBio lower CI ≥ 0.05 threshold, 14 not easy-to-map, non-homopolymer variants passed, and totaled 135 mosaic database variants for manual curation (**Supplemental Figure 3: green boxes**). Variants that did not adhere to decision tree heuristics were excluded from the benchmark VCF. if they were likely false positives or <2% VAF. Variants were excluded from both VCF and BED files if the evidence was unclear or the VAF could not be confidently determined to be < 0.02 or ≥ 0.05 (**Supplemental Figure 3: red, yellow boxes**).

### Mosaic Benchmark set Characterization and External Validation

HG002 mosaic benchmark variants in medically relevant genes were identified by intersecting the benchmark VCF with a BED file containing coordinates for genes in a previously generated curated list of 5,026 medical relevant genes (MRGs) (Wagner, Olson, Harris, McDaniel, et al. 2022). MRG coverage info and the number of bases occurring in HG002 mosaic benchmark regions were obtained by comparing the mosaic benchmark BED and GRCh38 full MRG BED files.

The draft mosaic benchmark set was shared with eight groups working on somatic or mosaic variant calling methods for external validation. Each group compared their somatic or mosaic callset(s) against a draft version of the HG002 mosaic benchmark to determine if the benchmark set can be used to reliably identify errors (Olson et al. 2023). Tumor-normal and/or tumor-only mode(s) were used to generate callsets with commercial and open-source variant calling tools. Methods from the eight groups are available in the Supplemental Methods. Using the hap.py (Krusche, n.d.) benchmarking tool with vcfeval (Cleary et al. 2015) option enabled, the external callsets were compared against the HG002 mosaic benchmark v1.0 and also a combined HG002 mosaic benchmark v1.0 plus GIABv4.2.1 HG002 germline small variant benchmark VCF to assess if the benchmark could reliably identify false positives.

## Supplemental Information

See Supplemental Methods for external callset methods.

## Supplemental Figures

**Supplemental Figure 1:**
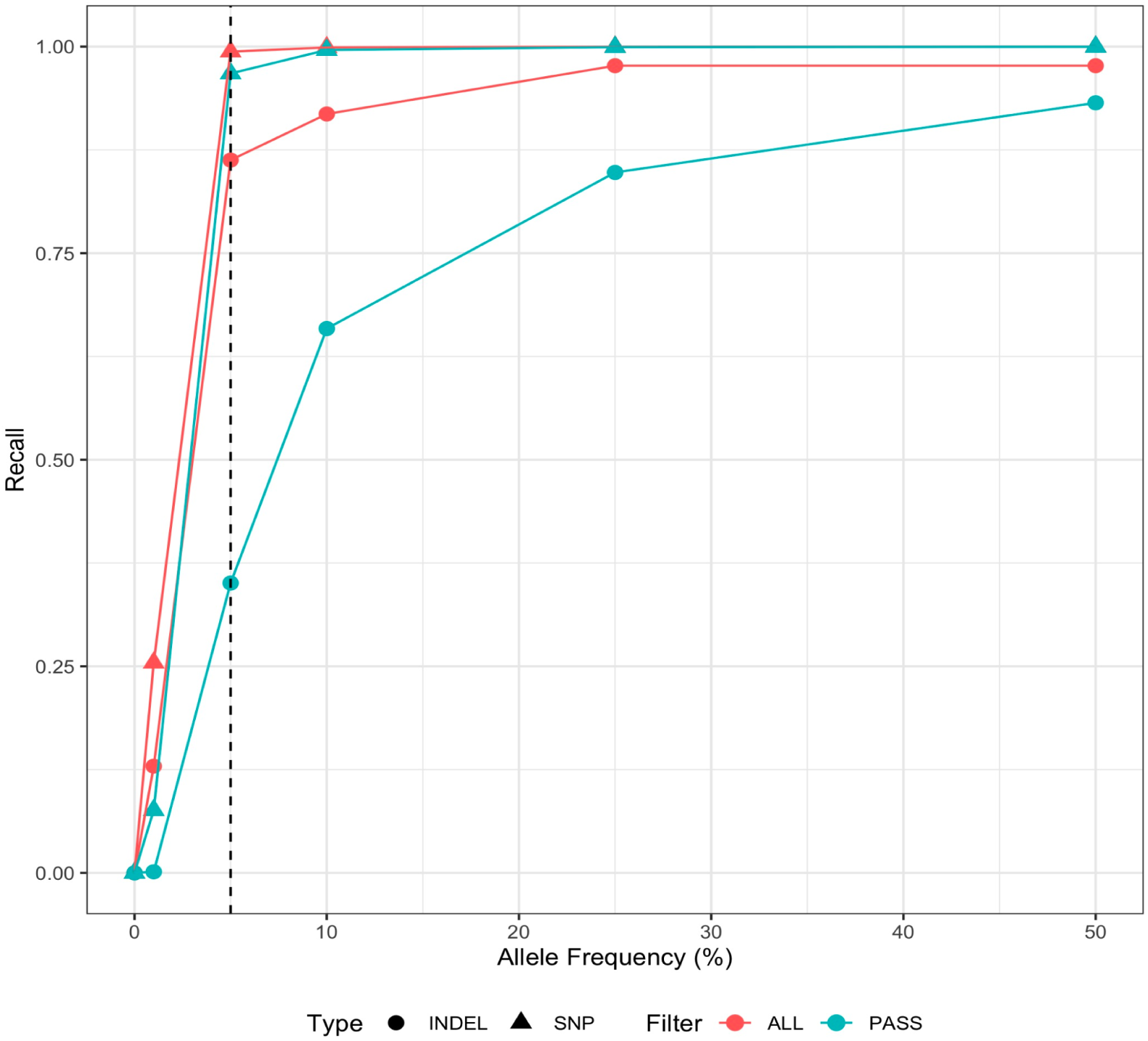
Limit of detection (LOD) was established at 5% variant allele fraction (VAF) using Strelka2 callsets from six in-silico mixtures of GIAB reference 300x samples (subset to chromosome 20), HG002 (son) and HG003 (father), as a control. X-axis values indicate the different samples, ranging from 0% AF (HG003 unmixed) to 50% AF (HG002 unmixed). Callsets for each mixture were benchmarked in the precisionFDA app.

**Supplemental Figure 2:**
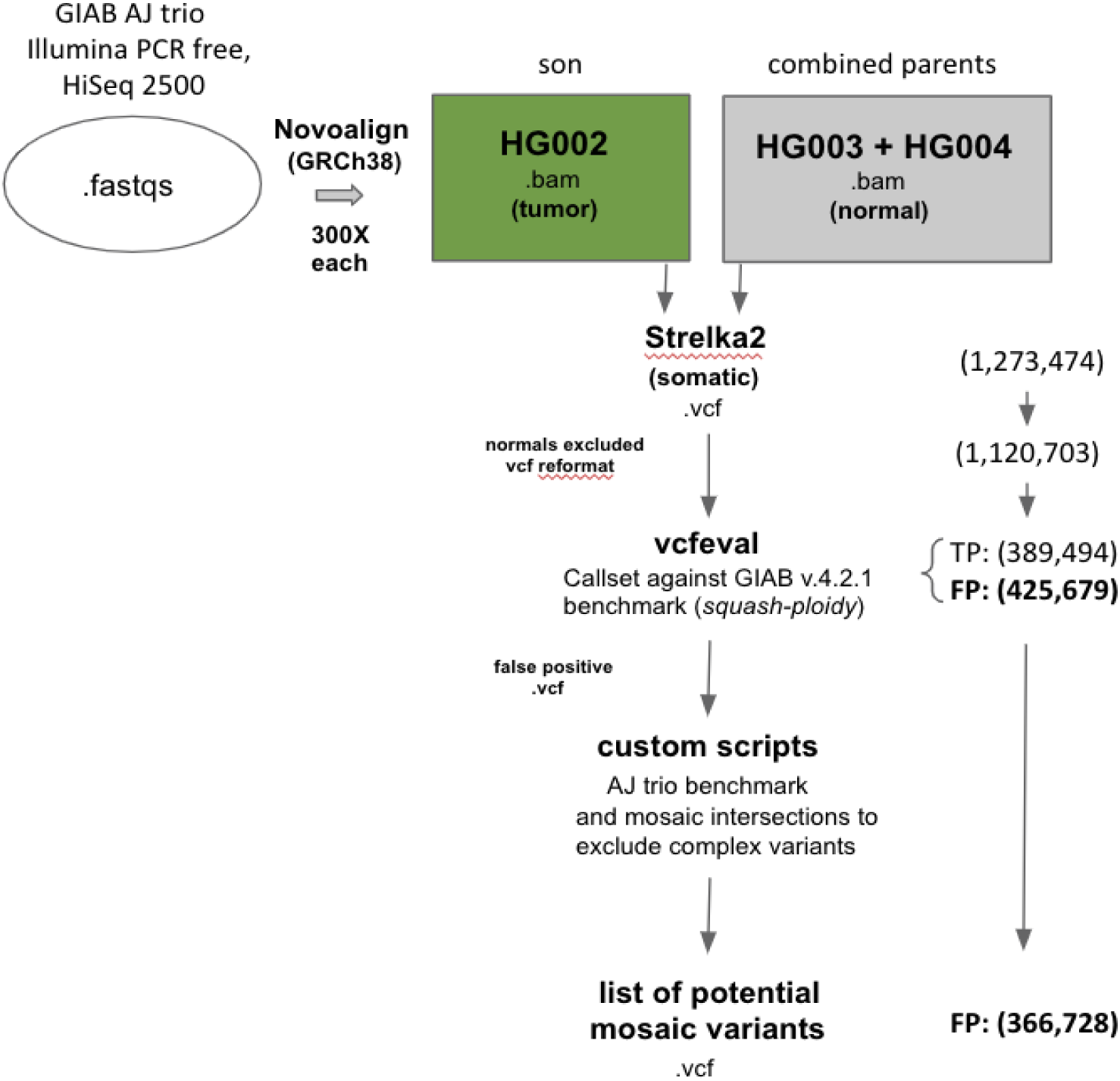
Variant counts from the Strelka2 tumor-normal run with the GIAB AJ trio, benchmarking, and filtering steps to generate a list of potential mosaic variants (366,728 vcfeval false positives) for database creation.

**Supplemental Figure 3:**
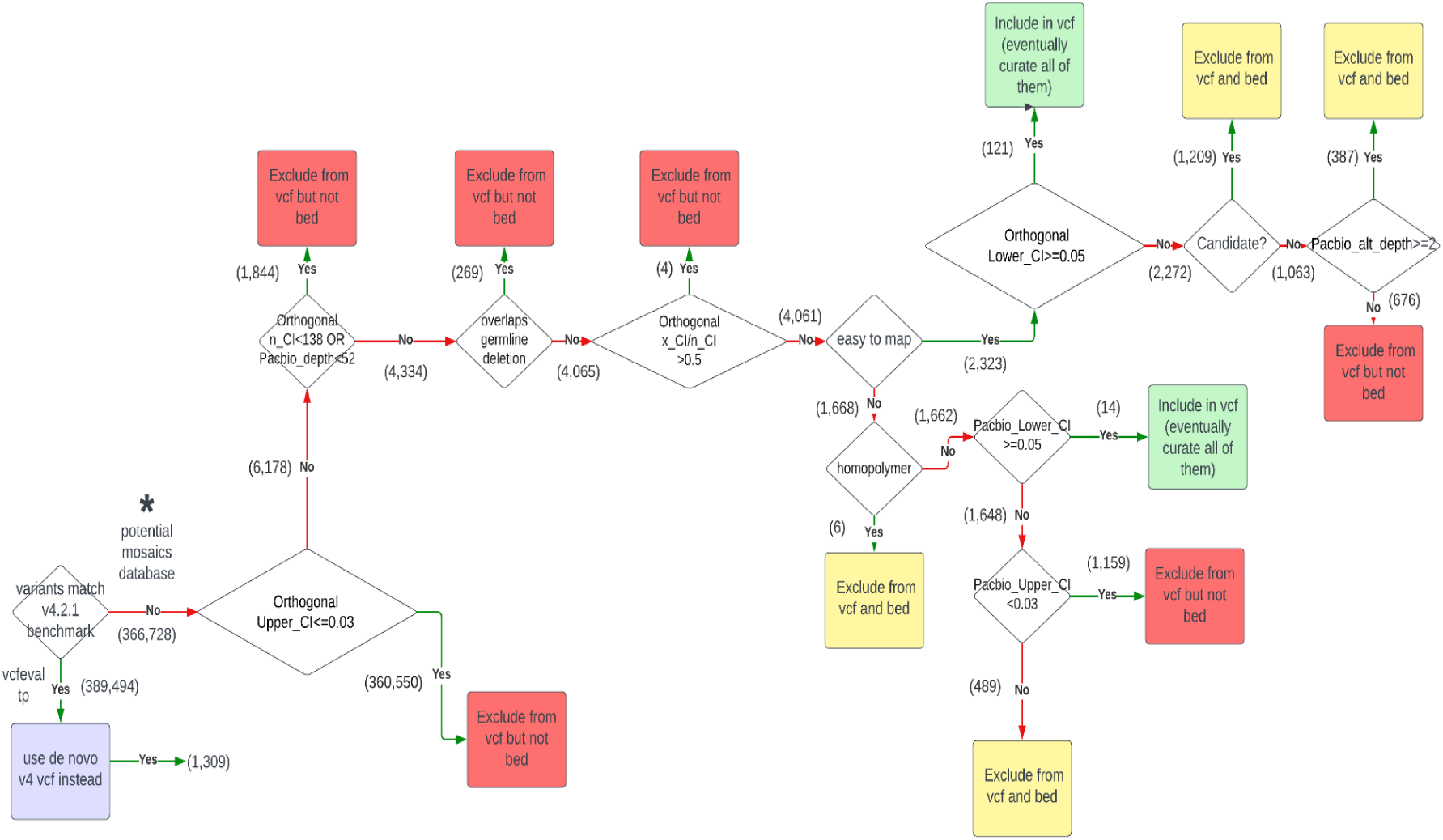
Mosaic benchmark decision tree for filtering potential mosaic variants to produce a list for manual curation. A series of heuristics were applied to the potential mosaic database (* starting at bottom left) using combined orthogonal CI thresholds and other attributes for candidate set determination. The tree resulted in three groups of variants: green boxes indicate variants to curate, yellow represents variants excluded from benchmark VCF and BED, and red variants excluded from benchmark VCF but not BED. Values in parentheses are variant counts for each step of the decision tree.

**Supplemental Figure 4:**
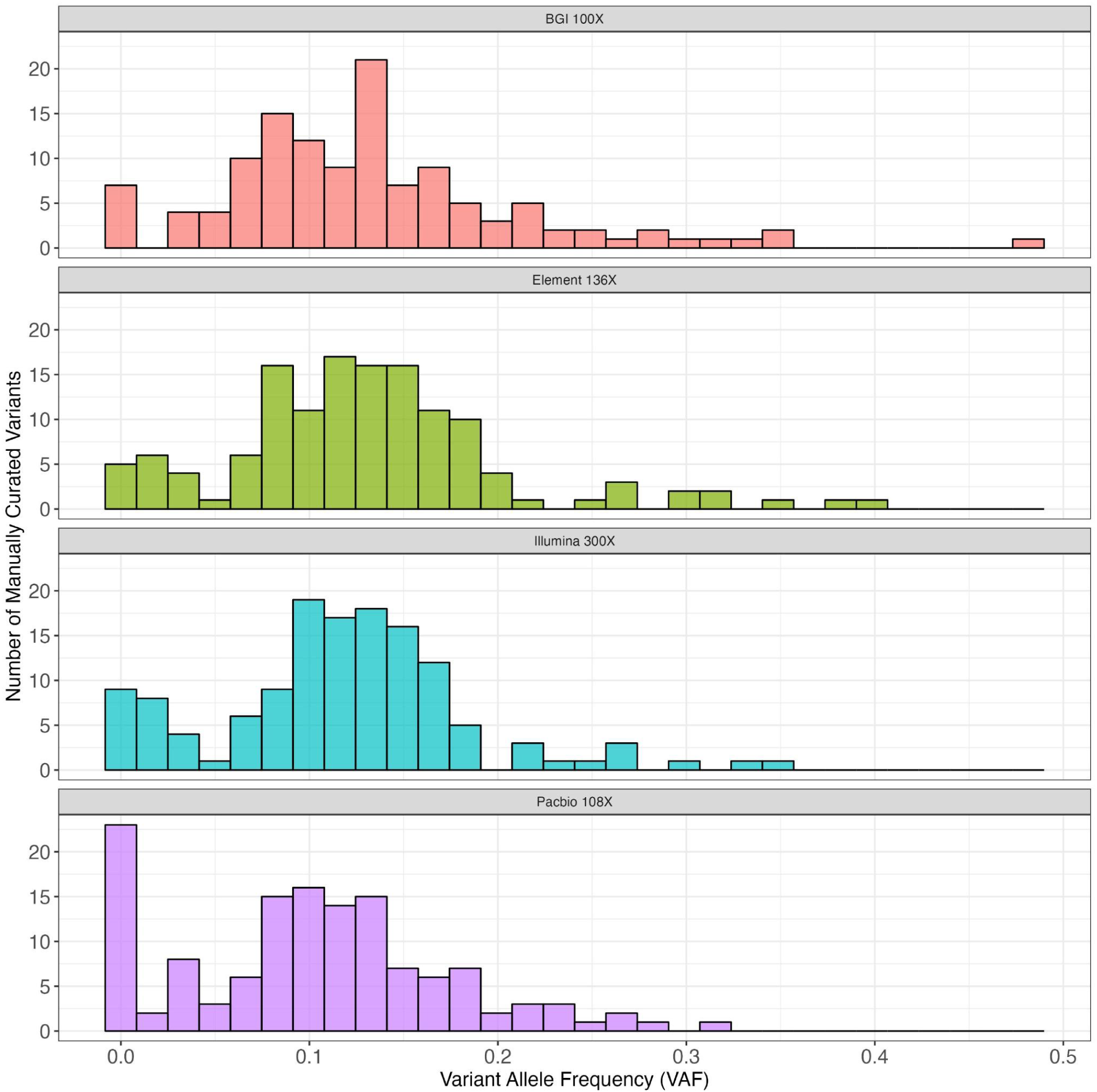
VAF distributions for all **<u>manually curated variants</u>** (135) in high-coverage Illumina (300x) and orthogonal (BGI - 100x, Element - 136x, PacBio HiFi - 106x) datasets generated from NIST HG002 reference material.

**Supplemental Figure 5:**
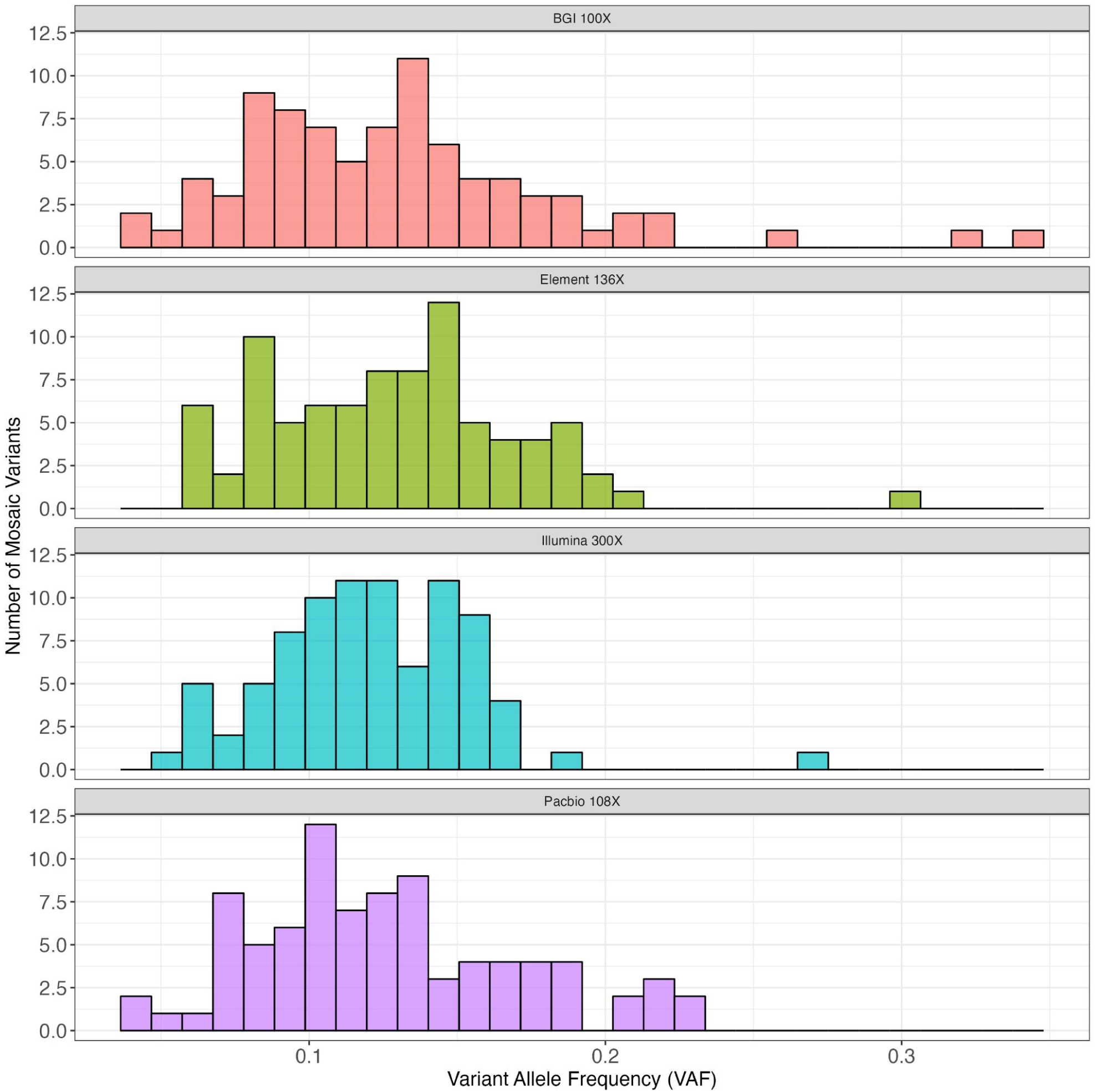
VAF distributions for the 85 HG002 **<u>mosaic benchmark variants</u>** in high-coverage Illumina (300x) and orthogonal (BGI - 100x, Element - 136x, PacBio HiFi - 106x) datasets generated from NIST HG002 reference material.

**Supplemental Figure 6:**
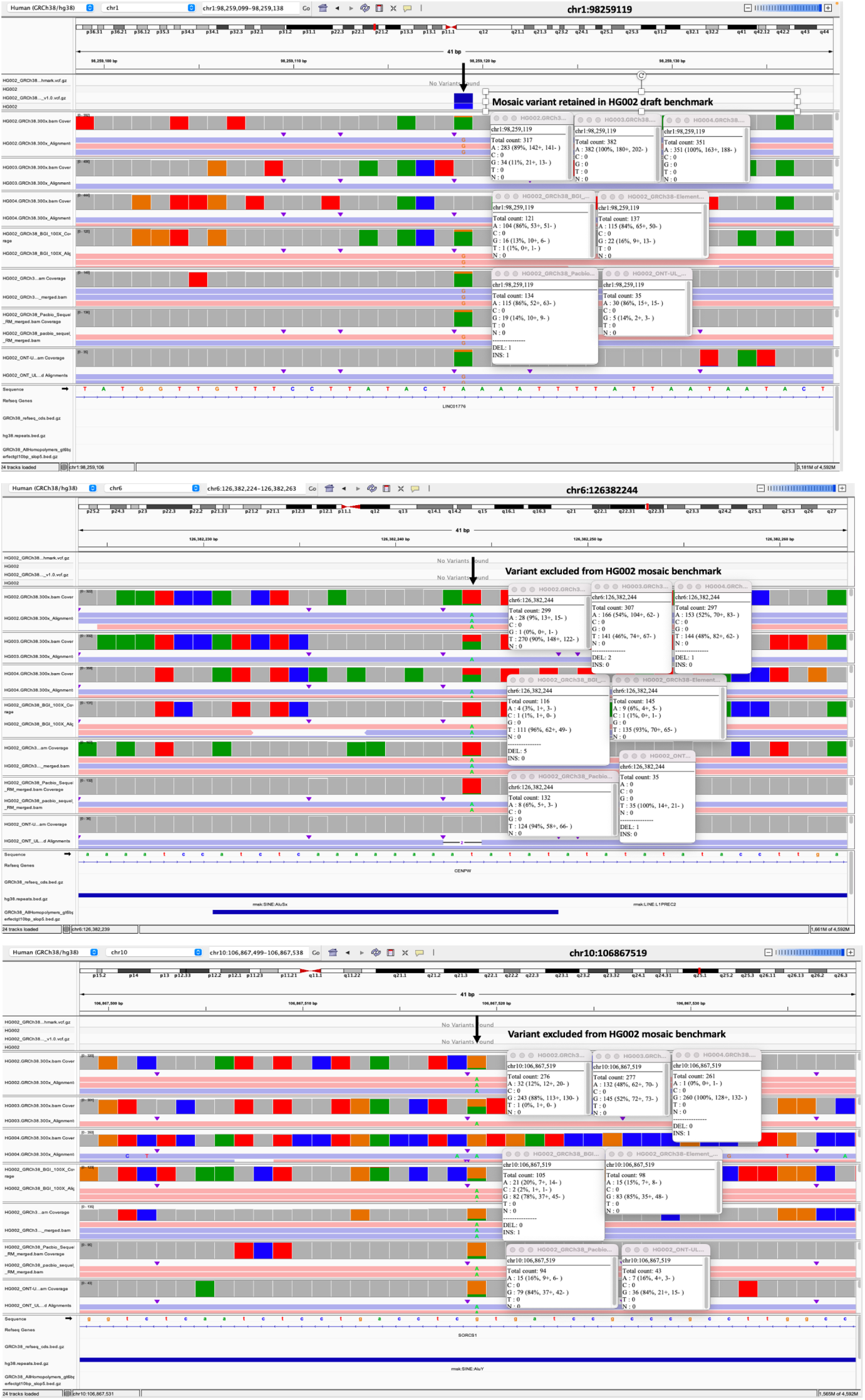
IGV profiles of variants (A) included in and (B,C) excluded from the HG002 mosaic benchmark after manual curation. Profile A depicts an HG002-specific candidate mosaic SNV (chr1:98259119 - not detected in parents), while B and C show a putative and candidate SNV (chr6:126382244 and chr10:106867519). The first three data tracks are the HG002 (son), HG003 (mother), and HG004 (father) 300X coverage Illumina data. The following tracks are for HG002 BGIseq dataset, Element, PacBio Sequel, and ONT-UL datasets.

## Data availability

The HG002 mosaic benchmark v1.1 files and external validation VCFscan be found at the Genome in a Bottle ftp site: https://ftp-trace.ncbi.nlm.nih.gov/ReferenceSamples/giab/release/AshkenazimTrio/HG002_NA24385_son/mosaic_v1.10/GRCh38/SNV

Mosaics database used to generate benchmark is located at https://ftp-trace.ncbi.nlm.nih.gov/ReferenceSamples/giab/release/AshkenazimTrio/HG002_NA24385_son/mosaic_v1.10/GRCh38/SNV/SupplementaryFiles/

Medically relevant gene (MRG) full BED from Wagner et al 2022 is here: https://github.com/usnistgov/cmrg-benchmarkset-manuscript/blob/master/data/gene_coords/unsorted/GRCh38_mrg_full_gene.bed

Sequencing data and data resources used to generate and characterize the mosaic benchmark set are listed in Supplemental Table 1.

## Code availability

Code used in this study can be accessed at the following github repo: https://github.com/usnistgov/giab-HG002-mosaic-benchmark

## Author contributions

J.Z. and N.D.O. designed the study. J.Z., N.D.O., C.D., A.A., and M.H.C. performed analyses. D.J., L.A.R-R, A.M-B, J.M.L-S, C.F.B.Y., S.M.E.S, Y.W., M.R., A.V., L.M., W-T.C., S.C., J.H., R.M., G.P., A.C., P-C.C., K.S., D.C., A.K., L.B., M.F.E.M., Y.P., T.N.Y, P.C.B., K.S., J.F., C.E.M., B.R.L., C.A.R-R, and S.K. participated in the external evaluation of the draft benchmark set. C.D., N.D.O., A.A., M.H.C., M.M., and J.Z. wrote the manuscript. All authors reviewed and approved the manuscript.

## Acknowledgements

The authors would like to thank Kevin Keissler, Justin Wagner, and Maryellen de Mars for feedback on the draft manuscript. Certain equipment, instruments, software, or materials are identified in this paper in order to specify the experimental procedure adequately. Such identification is not intended to imply recommendation or endorsement of any product or service by NIST, nor is it intended to imply that the materials or equipment identified are necessarily the best available for the purpose.

## Funding

### Cornell

C.E.M. thanks the Scientific Computing Unit (SCU), the WorldQuant and GI Research Foundations, NASA (80NSSC23K0832, the National Institutes of Health (R01ES032638, U54AG089334), and the LLS (MCL7001-18, LLS 9238-16, 7029-23).

### ITER

This work was supported by Instituto de Salud Carlos III [CB06/06/1088 and PI23/00980],and co-financed by the European Regional Development Funds, “A way of making Europe” from the European Union; Ministerio de Ciencia e Innovación [RTC-2017-6471-1, AEI/FEDER, UE]; Cabildo Insular de Tenerife [CGIEU0000219140 and A0000014697]; by the agreement with Instituto Tecnológico y de Energías Renovables (ITER) [OA23/043] to strengthen scientific and technological education, training, research, development and scientific innovation in genomics, epidemiological surveillance based on massive sequencing, personalized medicine, and biotechnology; and by the agreement between Consejería de Educación, Formación Profesional, Actividad Física y Deportes and Cabildo Insular de Tenerife (AC0000022149).

### MDIC

This work, as part of the Somatic Reference Samples Initiative, is funded by grants from Illumina, Quidel, Gordon and Betty Moore Foundation and National Philanthropic Trust.

### UCLA

P.C.B., Y.P., T.N.Y., and M.F.E.M. were supported by the NIH through awards P30CA016042, U2CCA271894, U24CA248265, U54HG012517 and by the DOD PCRP award W81XWH2210247.

