## Supplemental Methods for "A robust benchmark for detecting low-frequency variants in the HG002 Genome In A Bottle NIST reference material"

Supplemental Methods: External Callsets

#### **Team Name**: Children's Mercy Kansas City

**Team Members:** Byunggil Yoo

##### General Description of Methods

**Input data**:
For this benchmark, we used Illumina PCR-free whole genome (WGS) data 2x150bp 40X per individual FASTQ files sequenced at Children's Mercy Kansas City.

**Read alignment**:

- DRAGEN 4.0.3 performed the alignment on the multi genome graph reference available here: https://support.illumina.com/downloads/dragen-reference-genomes-hg38.html
    - Alignments and Mark duplicates

```
dragen --config alignment.cfg
```

alignment.cfg includes

```
enable-map-align = true
enable-map-align-output = true
enable-bam-indexing = true
enable-sort = true
enable-deterministic-sort = true
enable-duplicate-marking = true
remove-duplicates = false

ref-dir = hg38+alt_masked+cnv+graph+hla+rna-8-r2.0-1
```

- **Tools:**
    - Illumina DRAGEN Bio-IT Platform v4.0.3.

**Mosaic variant calling**:

DRAGENE 4.0.3 somatic variant calling in tumor only mode detects the mosaic variant calls.

```
dragen --config somatic-variant-calling.cfg --tumor-bam-input alignment.bam
```

somatic-variant-calling.cfg includes

```
enable-map-align = false
enable-map-align-output = false
enable-bam-indexing = false
enable-sort = true
enable-deterministic-sort = true
enable-duplicate-marking = false
remove-duplicates = false
ref-dir = hg38+alt_masked+cnv+graph+hla+rna-8-r2.0-1
enable-variant-caller = true
vc-hard-filter = DRAGENHardSNP:snp: MQ < 30.0 || MQRankSum < -12.5 ||
ReadPosRankSum < -8.0;DRAGENHardINDEL:indel: ReadPosRankSum < -20.0
vc-max-alternate-alleles = 6
vc-target-coverage = 2000
vc-min-read-qual = 20
vc-min-base-qual = 10
vc-min-call-qual = 20.0
vc-min-reads-per-start-pos = 5
vc-emit-zero-coverage-intervals = true
vc-decoy-contigs = chrEBV
vc-decoy-contigs = hs38d1
enable-smn = false
enable-cyp2d6 = false
enable-hrd = false
vc-ml-enable-recalibration = true
```

- **Tools:**
    - Illumina DRAGEN Bio-IT Platform v4.0.3.

**Benchmark curation decision**:

- Finally, using the **"**MOSAIC >5% VAF?" column from the *mosaic benchmark curation sheet*, we declared:
    - If our procedure detected the variant, our verdict was TRUE, because our data is relatively low depth and MOSAIC >5% VAF is required to be called.
    - If our procedure didn't detect the variant, our verdict was FALSE
    - We performed a manual curation with IGV to complement our verdict in the cases of discordance between our verdict and the GIAB call. The manual curation is left on the column, "NOTES"

### **Team Name**: Cornell (Mason Lab)

**Team Members:** Karolina Sienkiewicz, Jonathan Foox, Christopher E Mason

**General Description of Methods**

**Input data**:

The Illumina PCR-free WGS( 2x150bp, 300X) FASTQ files for sample HG002 were retrieved based on indexes from NIST's GIAB GitHub repository. The reference genome file GRCh38-GIABv3 version was retrieved from the GIAB FTP repository. A set of indexes was created utilizing bwa-mem2, GATK CreateSequenceDictionary, and samtools. The known SNP and indels references for GRCh38 assembly were downloaded from the GATK Public Resource Bundle.

**Read alignment and mapping**:

- Each replicate from each run was processed using Sentieon's TNscope DNAseq workflow with the following steps:
    - Read alignments

```
readgroup=@RG\\tID:${sample}\\tSM:${sample}\\tPL:ILLUMINA
reference=GRCh38_GIABv3_no_alt_analysis_set_maskedGRC_decoys_MAP2K3_KMT2C_KCNJ18.f
    asta sentieon bwa mem -M -R "${readgroup}" -t $CORES -K 10000000 $reference $fastqR1
        $fastqR2 | sentieon util sort -r $reference -o ${sample}.bam -t $CORES --sam2bam -i -
```

- Collect alignment metrics

```
bam=${sample}.bam
sentieon driver -r $reference -t $CORES -i $bam \
        --algo MeanQualityByCycle MQmetrics_$sample.txt \
```

```
        --algo QualDistribution QDmetrics_$sample.txt \
        --algo GCBias --summary GCsummary_$sample.txt GCmetrics_$sample.txt \
        --algo AlignmentStat ALNmetrics_$sample.txt \
        --algo InsertSizeMetricAlgo ISmetrics_$sample.txt
sentieon plot GCBias -o GC_$sample.pdf GCmetrics_$sample.txt
sentieon plot QualDistribution -o QD_$sample.pdf QDmetrics_$sample.txt
sentieon plot MeanQualityByCycle -o MQ_$sample.pdf
MQmetrics_$sample.txt sentieon plot InsertSizeMetricAlgo -o
IS_$sample.pdf ISmetrics_$sample.txt
```

  - Mark and remove sequencing duplicates, followed by updating read tags

```
sentieon driver -t $CORES -i $bam --algo LocusCollector --fun score_info
${sample}_score.txt sentieon driver -t $CORES -i $bam --algo Dedup --score_info
${sample}_score.txt \
        --metrics DEDUPmetrics_$sample.txt ${sample}.md.bam
sentieon driver -r $reference -t $CORES -i ${sample}.md.bam \
        --algo CoverageMetrics DEDUPcovmetrics_$sample
```

  - Base quality score recalibration

```
known_snps=Homo_sapiens_assembly38.dbsnp138.vcf.gz
known_indels=Mills_and_1000G_gold_standard.indels.hg38.vcf.gz
sentieon driver -r $reference -t $CORES -i ${sample}.merged.bam --algo QualCal -k
$known_snps -k $known_indels RECAL_$sample.table
    sentieon driver -r $reference -t $CORES -i ${sample}.merged.bam -q
   RECAL_$sample.table \ --algo QualCal -k $known_snps -k $known_indels
                RECAL_$sample.table.post
sentieon driver -t $CORES --algo QualCal --plot --before RECAL_$sample.table \
        --after RECAL_$sample.table.post RECALdiff_$sample.csv
sentieon plot QualCal -o RECAL_$sample.pdf RECALdiff_$sample.csv
```

- Final BAM files per run for each sample were merged using bamtools and indexed using samtools

```
ls *md.bam > bam_list.txt
bamtools merge -list bam_list.txt -out HG002.merged.bam
samtools index -@ $CORES HG002.merged.bam HG002.merged.bam.bai
```

- Update the read tags in the merged BAM file

```
samtools view -H HG002.merged.bam | grep -v '@RG' > header.sam
samtools view -H HG002.merged.bam | grep '^@RG' | sed -e "s/SM:2/SM:HG002--/" | awk
-F'--' '{print $1}' >> header.sam
samtools reheader header.sam HG002.merged.bam > HG002.merged.rh.bam
```

- Tools:

        - Sentieon (v202010)
        - bamtools (v2.5.2)
        - samtools (v1.9)

**Mosaic variant calling**:

- Variant calling with Sentieon TNScope:

```
sentieon driver -r $reference -t $CORES -i HG002.merged.rh.bam -q
RECAL_HG002.table \ --algo TNscope --disable_detector sv --trim_soft_clip
--tumor_sample “HG002” \ -q RECAL_HG002.table --dbsnp $known_snps
HG002.somatic.vcf.gz
```

- **Tools:**
    - Sentieon (v202010)

**Benchmark curation decision**:

- Finally, using the KEEP/?/REMOVE column from the *mosaic benchmark curation sheet*, we declared:
    - If our procedure defined a variant as TP and it was marked as KEEP, our verdict was to KEEP it.
    - If our procedure defined a variant as FP and it was marked as REMOVE, our verdict was to REMOVE it.
    - Otherwise, we perform a manual curation with IGV to complement our verdict in the cases of discordance between our verdict and the GIAB call.

### Team Name: DRAGEN (Illumina)

**Team Members:**
Yina Wang, Massimiliano Rossi, Arun Visvanath, Lisa Murray, Wei-Ting Chen, Severine Catreux, James Han, Rami Mehio, Gavin Parnaby

#### General Description of Methods

In DRAGEN v4.3 we added mosaic detection within the DRAGEN germline pipeline, using an advanced machine learning (ML) model to detect SNP and indel mosaic variants. Our mosaic detection algorithm exploits the DRAGEN pangenome reference to recover low allele frequency (AF) calls, without requiring matched controls. Mosaic calls are integrated into a standard VCF output alongside germline variants, using tags to ease interpretation.

In Fig. 1, we present both the default DRAGEN-ML germline workflow (A) and the enhanced DRAGEN-ML workflow with mosaic variant detection enabled (B). Users can enable/disable mosaic detection in the germline workflow as desired. Our mosaic detection workflow achieves remarkable recall and accuracy through three key enhancements: (1) We improve sensitivity by recovering reads in low-mappability regions using the DRAGEN pangenome reference and associated advanced alignment algorithms (2) We extract active regions with a lower evidence threshold. This allows positions with lower read evidence to progress through the pipeline increasing sensitivity. (3) We use a ML model that is trained specifically to identify mosaic variants improving specificity. The model runs after the germline pipeline has identified putative germline variants and identifies lower-AF mosaic calls in the remaining variant candidates.

The mosaic model is trained using supervised learning. Due to a shortage of authentic & validated mosaic data, we use Bamsurgeon to simulate mosaic variants (both SNP and INDEL). ~50k and ~10k mosaic variants are generated in GIAB v4.2.1 truth bed reference-homozygous positions for WGS and WES data respectively. The mosaic variant AF follows a uniform distribution ranging from 1% to 45%. We simulate mosaic variants in a range of sequencing platforms and configurations so that the model generalizes well across different sequencers, depths, lab-preparation flows, coverages, etc. We test our model using real mosaic data, admixture datasets, and reference datasets.

We train the mosaic model using rich read level features including statistical descriptions of mapping quality, base quality, strand bias, variant length, GC bias, depth, AF as well as internal HMM scores including foreign read probabilities, SSE triggers, base quality, and other statistics from VC internal processing. These features are extracted during DRAGEN variant calling. The features are used to build a model using offline training, outside the DRAGEN pipeline. The model uses a gradient-boosted ensemble of weak decision tree learners to identify mosaic variants, resulting in a very efficient and accurate model (adds only a few minutes to variant calling time without requiring hardware acceleration).

Mosaic variants are output in the same VCF file as germline variants. For a called mosaic variant, we tag the record's INFO field using a MOSAIC tag, and we set genotype (GT) to 0/1. We update the QUAL field with a confidence score calculated from the model probability output. We calibrated the mosaic pass threshold on the QUAL field to recover high confidence mosaic events based on validated mosaic data.

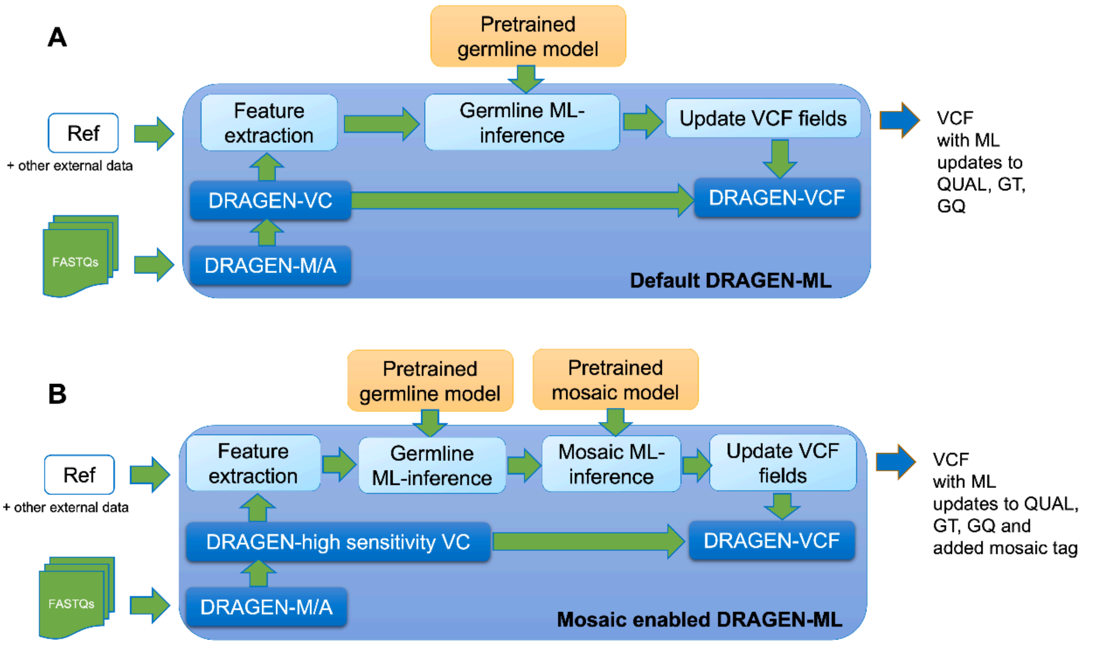


*Figure 1: (A) Default DRAGEN ML workflow; (B) Mosaic enabled DRAGEN-ML workflow.*

**Input data**:
For this benchmark evaluation, we used the GIAB HG002 Illumina PCR-free WGS (2x150bp, 300X) FASTQs.

**End-to-end mosaic variant calling**:
- Variant calling tool(s) – DRAGEN v.4.3.6, end-to-end DNA pipeline run, high-sensitivity mode, multigenome reference enabled

```
dragen \
--fastq-list=<path-to-hg002_300x_fastq-list> \
--ref-dir=<path-to-hg38-alt_masked.graph.cnv.hla.rna_v3> \
```

```
--output-file-prefix=HG002_HiSeq_300x \
--output-directory=<path-to-output-directory> \
--events-log-file dragen_events.csv \
--vc-enable-mosaic-detection=true \
--generate-sa-tags=true \
--enable-vcf-compression=true \
--enable-variant-caller=true \
--enable-map-align=true \
--enable-map-align-output=true \
--enable-sort=true \
--enable-duplicate-marking=true \
--enable-bam-indexing=true
```

**Mosaic variant filtering**:

- Mosaic variants are tagged with MOSAIC vcf INFO tag and can be filtered with bcftools.
- We used bcftools to generate a mosaic-only hard-filtered VCF file using the following command:

bcftools filter -i "(INFO/MOSAIC==1)" HG002_300x.hard-filtered.vcf.gz -Oz > HG002_300x.mosaic-only.hard-filtered.vcf.gz

### **Team Name**: Element Biosciences

**Team Members:** Bryan R. Lajoie, Carlos Ruiz, Mitch Sudkamp, Mark Ambroso, Shawn Levy, Semyon Kruglyak

#### General Description of Methods

**Input data**:

For this benchmark evaluation, we used the GIAB AJ trio (HG002/HG003/HG004) NIST RM and sequenced using our new Element UltraQ (Q50) chemistry. The VCF was derived from 70x tumor/normal data, with 70x HG002 as the "tumor" and a 70x HG003+HG004 synthetic mix as the "normal".

**Read alignment and mapping**:

- Each sample was processed using the following steps:
    - **Alignment** -Sentieon BWA (Sentieon-v2023.08.0) aligned to the standard hg38 (Homo_sapiens_assembly38) reference, and using the DNAscopeElementBioWGS2.0.bundle/bwa.model model.

```
sentieon bwa mem \
    -x DNAscopeElementBioWGS2.0.bundle/bwa.model \
    -M -R "@RG\tID:MAXQ-0216__GAT-APP-C138\tSM:GAT-APP-C138\tPL:ELEMENT" \
    -t 68 \
    -K 10000000 \
    $INDEX \
    GAT-APP-C138_FQD-2x150x150-70x_R1.fastq.gz GAT-APP-C138_FQD-2x150x150-70x_R2.fastq.gz \
    | sentieon util sort $bam_option -r Homo_sapiens_assembly38.fa -o GAT-APP-C138__MAXQ-0216.bam -t
68 --sam2bam -i -
```

- Duplicates - marking and removal via Sentieon LocusCollector & Dedup (Sentieon-v2023.08.0)

```
sentieon driver \
    -t 24 \
    -i GAT-APP-C138__MAXQ-0216.bam \
    --algo LocusCollector \
    --fun score_info \
    GAT-APP-C138.score.txt

 sentieon driver \
        -t 24 \
        -i GAT-APP-C138__MAXQ-0216.bam \
        --algo Dedup \
        --rmdup \
        --score_info GAT-APP-C138.score.txt \
        --optical_dup_pix_dist 100 \
        --metrics GAT-APP-C138.dedup_metrics.txt \
        GAT-APP-C138.deduped.bam
```

**Mosaic variant calling**:

- Variant Calling - Google Deepsomatic v1.6.1 (docker.io/google/deepsomatic:1.6.1). A 70x downsampling of HG002 was used as the “tumor reads”. A 70x downsampling of a synthetic HG003/HG004 was used as the “normal reads”. To generate the 70x HG003/HG004, we synthetically merged a 35x HG003 and a 35x HG004 from two existing runs (MAXQ-0188 and MAXQ-0189).

```
run_deepsomatic \
    --model_type WGS \
    --ref Homo_sapiens_assembly38.fa \
    --reads_normal=GAT-APP-C140__GAT-APP-C142.deduped.bam \
    --reads_tumor=GAT-APP-C138.deduped.bam \
    --output_vcf=GAT-APP-C138.deepsomatic.output.vcf.gz \
    --sample_name_tumor=GAT-APP-C138 \
    --sample_name_normal=GAT-APP-C140__GAT-APP-C142 \
    --num_shards 68 \
    --intermediate_results_dir /tmp/intermediate_results_dir
```

**Benchmark curation decision**:

- Finally, using the KEEP/?/REMOVE column from the *mosaic benchmark curation sheet*, we declared:
    - If our procedure defined a variant as TP and it was marked as TP, our verdict was to KEEP it.
    - If our procedure defined a variant as FP and it was marked as FP, our verdict was to REMOVE it.
    - Otherwise, we perform a manual curation with IGV to complement our verdict in the cases of discordance between our verdict and the GIAB call.

**AVITI UltraQ 70x**

**422 total calls**

**85/85 detected (any filter)**

**81/85 TP (PASS)**

**4/85 (FN, labeled RefCall via DS)**

0/85 absent

### **Team Name**: Google Research Genomics Team

**Team Members:** Andrew Carroll, Pi-Chuan Chang, Kishwar Shafin, Daniel Cook, Alexey Kolesnikov, Lucas Brambrink

#### General Description of Methods

**Input data**:

Revio 130x data taken directly from GIAB IGV session
Illumina 300x data taken directly from GIAB IGV session
Element Cloudbreak data (~100x) from
https://www.biorxiv.org/content/10.1101/2023.08.11.553043v1
(https://storage.mtls.cloud.google.com/brain-genomics-public/research/element/cloudbreak_wgs/HG002.element.cloudbreak.500bp_ins.grch38.bam)
Onso data from PacBio downloads
(https://downloads.pacbcloud.com/public/onso/2023Q3/WGS/hg002_30x_WGS/)

**Read alignment and mapping**:

- Each replicate from each run was processed using the following steps:
  - Alignments

```
BWA MEM (short reads)
Minimap2 (long reads)
```

  - Mark duplicates.

No duplicate marking (DeepVariant benchmarks ambivalent to whether marked or not)

  - Base quality score recalibration.

No BQSR (benchmarks indicate BQSR is slightly destructive to information content and

- Replicates per run from the same sample were merged using MergeSamFiles from GATK. Final BAM files per run for each sample were merged using MarkDuplicates from GATK.

No merging necessary here. BAM files taken directly from GIAB were already merged, or in case of Element and Onso came directly from FASTQ

- **Tools:**
  - BWA-MEM and DeepSomatic

**Mosaic variant calling**:
- Tumor-normal or tumor-only calling:
  - DeepSomatic - v1.6 out-of-the box model. No retraining or custom models

```
 Command used:

#Illumina, Element, Onso
INPUT_DIR=${PWD}/input
OUTPUT_DIR=${PWD}/output
BIN_VERSION=1.6.0

sudo docker run \
     -v ${INPUT_DIR}:${INPUT_DIR}/ \
     -v ${OUTPUT_DIR}:${OUTPUT_DIR}/ \
     google/deepsomatic:"${BIN_VERSION}" \
     run_deepsomatic \
     --model_type=WGS \
     --ref=${INPUT_DIR}/GRCh38.no_alt_analysis_set.fa.gz \
     --reads_normal=${INPUT_DIR}/${NORMAL} \
     --reads_tumor=${INPUT_DIR}/${TUMOR} \
     --output_vcf=${OUTPUT_DIR}/${VCF} \
     --sample_name_tumor="tumor" \
     --sample_name_normal="normal" \
     --num_shards=$(nproc)

#PacBio

INPUT_DIR=${PWD}/input
OUTPUT_DIR=${PWD}/output
BIN_VERSION=1.6.0

sudo docker run \
     -v ${INPUT_DIR}:${INPUT_DIR}/ \
     -v ${OUTPUT_DIR}:${OUTPUT_DIR}/ \
```

```
    google/deepsomatic:"${BIN_VERSION}" \
    run_deepsomatic \
    --model_type=PACBIO \
    --ref=${INPUT_DIR}/GRCh38.no_alt_analysis_set.fa.gz \
    --reads_normal=${INPUT_DIR}/${NORMAL} \
    --reads_tumor=${INPUT_DIR}/${TUMOR} \
    --output_vcf=${OUTPUT_DIR}/${VCF} \
    --sample_name_tumor="tumor" \
    --sample_name_normal="normal" \
    --num_shards=$(nproc)
```

**Normal files:**

In each case, normal files were a ~30x mix of HG003 and HG004. In the case of short read variant calling (Illumina, Element, Onso) the normal used was NovaSeq (due to my judgment that it wouldn't matter much and because we don't have HG003/4 for Onso). The normal files used are publicly downloadable by these links:

https://storage.googleapis.com/brain-genomics/awcarroll/giab/mosaic/bams/HG003-HG004.normal.novaseq.grch38.bam
https://storage.googleapis.com/brain-genomics/awcarroll/giab/mosaic/bams/HG003-HG004.normal.novaseq.grch38.bam.bai
https://storage.googleapis.com/brain-genomics/awcarroll/giab/mosaic/bams/HG003-HG004.normal.pacbio.grch38.bam
https://storage.googleapis.com/brain-genomics/awcarroll/giab/mosaic/bams/HG003-HG004.normal.pacbio.grch38.bam.bai

- **Tools:**
    - DeepSomatic. https://github.com/google/deepsomatic

**Benchmark curation decision**:

- Finally, using the KEEP/?/REMOVE column from the *mosaic benchmark curation sheet*, we declared:
    - If our procedure defined a variant as TP and it was marked as KEEP, our verdict was to KEEP it.
    - If our procedure defined a variant as FP and it was marked as REMOVE, our verdict was to REMOVE it.
    - Otherwise, we perform a manual curation with IGV to complement our verdict in the cases of discordance between our verdict and the GIAB call.

DeepSomatic calls all 73 mosaic variants identified in the truth set. This is consistent across multiple technologies: Illumina, Onso, PacBio Revio. Element recover 71 of 73.

In addition Revio + one other short-read method (and usually all methods) call the following additional variants with >Q10 confidence (entries taken from Revio BAM).

chr1 106002189 . T G 29 PASS . GT:GQ:DP:AD:VAF:PL
0/1:29:159:126,33:0.207547:29,58,0
chr1 107008885 . T A 22.1 PASS . GT:GQ:DP:AD:VAF:PL
0/1:22:140:102,36:0.257143:22,42,0
chr2 178057258 . A G 29.9 PASS . GT:GQ:DP:AD:VAF:PL
0/1:30:157:115,42:0.267516:29,63,0
**chr5 20909983** . A G 12.4 PASS . GT:GQ:DP:AD:VAF:PL
0/1:12:171:144,27:0.157895:12,35,0
**chr6 150458314** . G T 23.6 PASS . GT:GQ:DP:AD:VAF:PL
0/1:24:130:101,29:0.223077:23,49,0
chr7 550099 . G A 30.2 PASS . GT:GQ:DP:AD:VAF:PL
0/1:30:92:70,22:0.23913:30,54,0
chr7 98565112 . G T 39.8 PASS . GT:GQ:DP:AD:VAF:PL
0/1:40:84:53,31:0.369048:39,64,0
chr10 18475692 . A T 15.4 PASS . GT:GQ:DP:AD:VAF:PL
0/1:15:108:91,17:0.157407:15,36,0
chr10 82665456 . C G 24.8 PASS . GT:GQ:DP:AD:VAF:PL
0/1:25:112:78,28:0.25:24,51,0
chr11 24797065 . C A 13.7 PASS . GT:GQ:DP:AD:VAF:PL
0/1:14:150:139,11:0.0733333:13,36,0
chr12 131194773 . G A 30.8 PASS . GT:GQ:DP:AD:VAF:PL
0/1:31:123:89,34:0.276423:30,54,0
chr13 26951923 . GA G 11.2 PASS . GT:GQ:DP:AD:VAF:PL
0/1:10:117:51,63:0.538462:10,15,0
chr13 27946048 . C T 15.9 PASS . GT:GQ:DP:AD:VAF:PL
0/1:16:111:93,16:0.144144:15,38,0
chr14 77871565 . C T 35.8 PASS . GT:GQ:DP:AD:VAF:PL
0/1:36:123:87,36:0.292683:35,56,0
chr20 677582 . A AAATAATAAT 18.8 PASS . GT:GQ:DP:AD:VAF:PL
0/1:18:109:45,56:0.513761:18,29,0

**Other notes on additional variants present -** The total number of somatic calls per sample were in the 1,000 - 10,000s. For each technology, I looked for whether there is a confidence cutoff that loses very little of the true calls. For the case on Onso, filtering doesn't seem to do all that much (and Onso is already the cleanest). For the others, Q10 looks like a good filter point. The table below covers the results:

| Dataset | Coverage | Quality Filter | Mosaic TPs | Additional SNP calls in Conf Region | Additional Indel calls in Conf Region |
|---|---|---|---|---|---|
| Onso | ~35x | 0 | 73 | 1975 | 703 |
| Revio | 130x | 0 | 73 | 1809 | 2445 |
| Illumina | 300x | 0 | 73 | 17545 | 5137 |
| Element | ~100x | 0 | 71 | 46783 | 3941 |
| Revio | 130x | 10 | 71 | 919 | 387 |
| Illumina | 300x | 10 | 71 | 5017 | 1103 |

| | | | | | |
|---|---|---|---|---|---|
| Element | ~100x | 10 | 69 | 5977 | 302 |

#### Team Name: Genomics Division at ITER

**Team Members:** David Jáspez; Luis Alberto Rubio-Rodríguez; Adrián Muñoz-Barrera; José Miguel Lorenzo-Salazar; Carlos Flores.

##### General Description of Methods

**Input data**:
For this benchmark, we used Illumina whole genome (WGS) data 2x150bp 300X per individual FASTQ files (HG002, HG003, and HG004 samples) downloaded from the GIAB repository (https://ftp-trace.ncbi.nlm.nih.gov/ReferenceSamples/giab/data_indexes/AshkenazimTrio/sequence.index.AJtrio_Illumina300X_wgs_07292015_updated).

**Read alignment and mapping**:

- Each replicate from each run was processed using the following steps from the GATK Best Practices guidelines:
    - BWA to generate alignments in SAM format using hg38 obtained from the GATK bundle.

```
ref="Homo_sapiens_assembly38.fasta"
bwa mem -K 100000000 -p -v 3 -t 16 -Y ${ref}
"HG002.run1.rep1.GRCh38.300x.unmapped.bam" | \
gatk MergeBamAlignment \
    -ALIGNED /dev/stdin \
    -UNMAPPED "HG002.run1.rep1.GRCh38.300x.unmapped.bam" \
    -O "HG002.run1.rep1.GRCh38.300x.BWA.bam" \
    -R ${ref} \
    -SO "unsorted" \
```

```
    --CREATE_INDEX true \
    --ADD_MATE_CIGAR true \
    --CLIP_ADAPTERS false \
    --CLIP_OVERLAPPING_READS true \
    --INCLUDE_SECONDARY_ALIGNMENTS true \
    --MAX_INSERTIONS_OR_DELETIONS -1 \
    --PRIMARY_ALIGNMENT_STRATEGY MostDistant \
    --ATTRIBUTES_TO_RETAIN XS \
    --VALIDATION_STRINGENCY SILENT \
    --EXPECTED_ORIENTATIONS FR \
    --MAX_RECORDS_IN_RAM 2000000 \
    --PROGRAM_RECORD_ID "bwamem" \
    --PROGRAM_GROUP_VERSION "0.7.17-r1188" \
    --PROGRAM_GROUP_COMMAND_LINE "-K 100000000 -p -v 3 -t 16 -Y ${ref}" \
    --PROGRAM_GROUP_NAME "bwamem" \
    --UNMAPPED_READ_STRATEGY COPY_TO_TAG \
    --ALIGNER_PROPER_PAIR_FLAGS true \
    --UNMAP_CONTAMINANT_READS true
```

- Mark duplicates.

```
gatk MarkDuplicates \
    -I "HG002.run1.rep1.GRCh38.300x.BWA.bam" \
    -O "HG002.run1.rep1.GRCh38.300x.BWA.deduped.bam" \
    -M "HG002.run1.rep1.GRCh38.300x.BWA.deduped.metrics" \
    --REMOVE_DUPLICATES false \
    --OPTICAL_DUPLICATE_PIXEL_DISTANCE 2500 \
    --VALIDATION_STRINGENCY SILENT \
    --ASSUME_SORT_ORDER queryname \
    --CREATE_MD5_FILE true \
    --CLEAR_DT false
```

- Base quality score recalibration (BQSR).

```
# Analyze patterns of covariation in the sequence dataset for BQSR
gatk BaseRecalibrator \
    -R ${ref} \
    -I "HG002.run1.rep1.GRCh38.300x.BWA.deduped.bam" \
    --use-original-qualities \
    -O "HG002.run1.rep1.GRCh38.300x.BWA.deduped.recal_data.table" \
    --known-sites "dbsnp_146.hg38.vcf" \
    --known-sites "Mills_and_1000G_gold_standard.indels.hg38.vcf"

# Apply the recalibration to your sequence data
gatk ApplyBQSR \
    -R ${ref} \
    -I "HG002.run1.rep1.GRCh38.300x.BWA.deduped.bam" \
    --use-original-qualities \
    --static-quantized-quals 10 \
    --static-quantized-quals 20 \
```

```
    --static-quantized-quals 30 \
    -bqsr "HG002.run1.rep1.GRCh38.300x.BWA.deduped.recal_data.table" \
    --create-output-bam-index \
    --create-output-bam-md5 \
    --add-output-sam-program-record \
    -O "HG002.run1.rep1.GRCh38.300x.BWA.deduped.recal.bam"
```

- Replicates per run from the same sample were merged using MergeSamFiles from GATK. Final BAM files per run for each sample were merged using MarkDuplicates from GATK.

```
gatk MergeSamFiles \
  -I "HG002.run1.rep1.GRCh38.300x.BWA.deduped.recal.bam" \
  -I "HG002.run1.rep2.GRCh38.300x.BWA.deduped.recal.bam" \
  -I "HG002.run1.rep3.GRCh38.300x.BWA.deduped.recal.bam" \
  -O "HG002.run1.merged.GRCh38.300x.BWA.deduped.recal.bam" \
  --ASSUME_SORTED true \
  --SORT_ORDER coordinate \
  --CREATE_INDEX true \
  --REFERENCE_SEQUENCE ${ref} \
  --VALIDATION_STRINGENCY SILENT

gatk MarkDuplicates \
  -I "HG002.run1.merged.GRCh38.300x.BWA.deduped.recal.bam" \
  -I "HG002.run2.merged.GRCh38.300x.BWA.deduped.recal.bam" \
  -O "HG002.GRCh38.300x.ITER.bam" \
  --CREATE_INDEX true \
  -M "HG002.GRCh38.300x.ITER.metrics" \
  --REMOVE_DUPLICATES false \
  --OPTICAL_DUPLICATE_PIXEL_DISTANCE 2500 \
  --VALIDATION_STRINGENCY SILENT \
  --ASSUME_SORT_ORDER coordinate \
  --CREATE_MD5_FILE true \
  --COMPRESSION_LEVEL 6
```

- **Tools:**
    - BWA, v0.7.15-r1188.
        - GitHub: https://github.com/lh3/bwa.
        - Li H. and Durbin R. Fast and accurate short read alignment with Burrows-Wheeler transform. Bioinformatics, 25, 1754-1760 (2009).
    - GATK4, v4.2.0.0.
        - https://gatk.broadinstitute.org/hc/en-us.
        - DePristo, M. A. et al. A framework for variation discovery and genotyping using next-generation DNA sequencing data. Nature Genet. 43, 491–498 (2011).

**Mosaic variant calling**:

- Tumor-only calling:
    - Mutect2 with default databases for germline resources and a panel of normals.

```
gatk Mutect2 \
```

```
  --reference ${ref} \
  --input "HG002.GRCh38.300x.ITER.bam" \
  --output "HG002.GRCh38.300x.ITER.Mutect2.vcf.gz" \
  --germline-resource "somatic-hg38_af-only-gnomad.hg38.vcf.gz" \
  --panel-of-normals "somatic-hg38_1000g_pon.hg38.vcf.gz" \
  --intervals "HG002_GRCh38_new_mosaic_benchmark_v0.1.bed" \
  --annotation MappingQualityRankSumTest \
  --annotation QualByDepth \
  --annotation ReadPosRankSumTest \
  --annotation RMSMappingQuality \
  --annotation FisherStrand \
  --annotation Coverage
```

- DRAGEN, in Somatic Mode with High Sensitivity Mode enabled and 'mean' prebuilt Systematic Noise Filtering in the regions of interest (ROI, defined as the set of candidate and putative variants plus a flanking region of 100 kb).

```
dragen \
 --force \
 --verbose \
 --ref-dir "/staging/references/hg38-alt_masked.cnv.hla.rna" \
 --tumor-bam-input "HG002.GRCh38.300x.ITER.ROI.bam" \
 --output-directory "${outdir}" \
 --output-file-prefix "HG002.GRCh38.300x.ITER.DRAGEN.vcf.gz" \
 --intermediate-results-dir "${tempdir}" \
 --enable-map-align false \
 --enable-sort false \
 --enable-metrics-json true \
 --enable-variant-caller true \
 --enable-vcf-compression true \
 --enable-vcf-indexing true \
 --vc-emit-ref-confidence BP_RESOLUTION \
 --vc-enable-vcf-output true \
 --vc-enable-high-sensitivity-mode true \
 --vc-target-bed "HG002_new_mosaic_benchmark_variants.padding100kb.bed" \
 --vc-systematic-noise
"/staging/resources/systematic_noise_files/systematic-noise-baseline-collection-
1.1.0/snv_wgs_hg38_mean_v1.1_systematic_noise.bed.gz" \
 --vc-enable-germline-tagging true \
 --enable-variant-annotation true \
 --variant-annotation-data "${nirvanadir}" \
 --variant-annotation-assembly GRCh38
```

- Tumor-normal calling:
  - Strelka2:
    - Using HG002 as tumor and HG003 as normal.
    - Using HG002 as tumor and HG004 as normal.

```
# Prepare Strelka2 script
${STRELKA_INSTALL_PATH}/bin/configureStrelkaSomaticWorkflow.py \
```

```
    --normalBam "HG003.GRCh38.300x.ITER.bam" \
    --tumorBam "HG002.GRCh38.300x.ITER.bam" \
    --referenceFasta ${ref} \
    --callRegions "HG002_GRCh38_new_mosaic_benchmark_v0.1.bed" \
    --runDir ${outdir}
# Run Strelka2 script
${outdir}/runWorkflow.py -m local -j 16
```

- RePlow:
  - Using HG002 as tumor and HG003 as normal.
  - Using HG002 as tumor and HG004 as normal.
  - And a series of complementary tests using technical replicates of HG002 and Somatic-only (Mutect2 call) and Somatic-and-Germinal model (Mutect2 and GATK4 calls, respectively).

```
java -jar ${replow} \
-r ${ref} \
-b "HG002.GRCh38.300x.ITER.ROI.bam"  \
-N "HG003.GRCh38.300x.ITER.ROI.bam" \
-T "HG002_new_mosaic_benchmark_variants.padding100kb.bed" \
-R ${Rscript} \
-o ${outdir} \
-L ${outfile_prefix}
```

- **Tools:**
  - Mutect2, GATK4 v4.2.0.0.
    - https://gatk.broadinstitute.org/hc/en-us.
    - DePristo, M. A. et al. A framework for variation discovery and genotyping using next-generation DNA sequencing data. Nature Genet. 43, 491–498 (2011).
  - Illumina DRAGEN Bio-IT Platform v4.2.4.
    - https://support-docs.illumina.com/SW/dragen_v42/Content/SW/FrontPages/DRAGEN.htm.
  - Strelka2, v2.9.10.
    - GitHub: https://github.com/Illumina/strelka.
    - Kim, S., Scheffler, K., Halpern, A.L. et al. Strelka2: fast and accurate calling of germline and somatic variants. Nat Methods 15, 591–594 (2018).
  - RePlow, v1.1.0.
    - https://sourceforge.net/projects/replow/.
    - Kim, J., Kim, D., Lim, J.S. et al. The use of technical replication for detection of low-level somatic mutations in next-generation sequencing. Nat Commun 10, 1047 (2019).

**Consensus calling and decision**:

- Preparation phase:
  - All variants from the *mosaic benchmark curation sheet* were extracted from each resulting VCF file for comparison.
  - Variants with a 'PASS' and VAF >5% were marked as TP.
  - Otherwise, FP.
- Discovery phase:

    - Obtain a consensus from the Strelka2, Mutect2, and DRAGEN calling results. Only if there was a call matching the three tools, the variant was marked as TP. Otherwise, the variant was considered a FP.
- Validation phase:
    - Relying on RePlow results:
        - If there was a TP match between the consensus of the discovery and validation phase, then the variant was marked as TP. Otherwise, the variant was considered a FP.
- Finally, using the KEEP/?/REMOVE column from the *mosaic benchmark curation sheet*, we declared:
    - If our procedure defined a variant as TP and it was marked as KEEP, our verdict was to KEEP it.
    - If our procedure defined a variant as FP and it was marked as REMOVE, our verdict was to REMOVE it.
    - Otherwise, we perform a manual curation with IGV to complement our verdict in the cases of discordance between our verdict and the GIAB call.

#### **Team Name**: NeuSomatic

**Team Members:** Sayed Mohammad Ebrahim Sahraeian, Roche Sequencing Solutions

##### General Description of Methods

NeuSomatic is an algorithm based on convolutional neural networks for accurate detection of somatic mutations. It can robustly detect somatic mutations across different sequencing platforms, strategies, and conditions, through proper training. NeuSomatic summarizes and augments sequence alignments in a unique manner, incorporating multi-dimensional features to effectively capture variant signals. In its ensemble mode, it can utilize information from other individual callers as additional input features to the network.

**Input data**:
For this benchmark, we used the GIAB AJ trio (HG002/HG003/HG004) Illumina PCR-free WGS (2x150bp, 300X) FASTQs.

**Read alignment and mapping**:

- Each replicate from each run was aligned as:

```
bwa mem -M
```

- Replicates per run from the same sample were merged using samtools merge.

- Mark duplicates for each sample

```
Picard MarkDuplicates I=input.bam O=output.dedup.bam
```

- HG003 and HG004 bams are also merged to form normal bam

- **Tools:**
  - BWA
    - GitHub: [https://github.com/lh3/bwa](https://github.com/lh3/bwa).
    - Li, H. (2013). [https://arxiv.org/abs/1303.3997](https://arxiv.org/abs/1303.3997).
    - Version: 0.7.15
  - Picard
    - webpage: https://broadinstitute.github.io/picard/
    - Version: 2.18.0

**Mosaic variant calling**:

- Tumor-normal calling:
  - NeuSomatic (ensemble mode using NeuSomatic_v0.1.4_ensemble_SEQC-WGS-GT50-SpikeWHGS10 model). In this model, in addition to the features/channels extracted from the tumor/normal bams, we used inputs from VarDict, MuTect2, Strelka2, MuSE, and SomaticSniper to define the set of input network channels for each candidate variants. The set of NeuSomatic calls where then detected as PASS SNV calls with AF in the range of 5-30% that overlap the high-confidence region of Mosaic variants on HG002.

```
neusomatic python preprocess.py --mode call --reference ref.fa --normal_bam
normal.bam --tumor_bam tumor.bam --work work --scan_maf 0.01 --min_mapq 10
--snp_min_af 0.01 --snp_min_bq 15 --snp_min_ao 2 --ins_min_af 0.02 --del_min_af
0.02 --ensemble_tsv ensemble_merged.tsv  --scan_window_size 100

python call.py --candidates_tsv work/dataset/*/c*tsv --reference ref.fa
--checkpoint NeuSomatic_v0.1.4_ensemble_SEQC-WGS-GT50-SpikeWGS10.pth --out work
--ensemble

python postprocess.py --reference ref.fa --tumor_bam .bam --pred_vcf
work/pred.vcf --candidates_vcf work/work_tumor/filtered_candidates.vcf
--output_vcf work/output.vcf --work work --ensemble_tsv ensemble_merged.
```

- **Tools:**
  - NeuSomatic
    - Github: [https://github.com/bioinform/neusomatic](https://github.com/bioinform/neusomatic)
    - Papers:
      [https://www.nature.com/articles/s41467-019-09027-x](https://www.nature.com/articles/s41467-019-09027-x)
      [https://genomebiology.biomedcentral.com/articles/10.1186/s13059-021-02592-9](https://genomebiology.biomedcentral.com/articles/10.1186/s13059-021-02592-9)
    - Version: 0.2.1
  - Individual callers used:
    - MuTect2 (4.4.0.0)
      - Cibulskis, K. et al. Sensitive detection of somatic point mutations in impure and heterogeneous cancer samples. Nat. Biotechnol. 31, 213 (2013).
    - SomaticSniper(1.0.5.0)
      - Larson, D. E. et al. SomaticSniper: identification of somatic point mutations in whole genome sequencing data. Bioinformatics 28, 311–317 (2011).
    - Strelka2 (2.9.5)

        - Kim, S. et al. Strelka2: fast and accurate calling of germline and somatic variants. Nat. Methods 15, 591–594 (2018).
    - MuSE (v1.0rc)
        - Fan, Y. et al. MuSE: accounting for tumor heterogeneity using a sample-specific error model improves sensitivity and specificity in mutation calling from sequencing data. Genome Biol. 17, 178 (2016).
    - VarDict (v1.5.1)
        - Lai, Z. et al. VarDict: a novel and versatile variant caller for next-generation sequencing in cancer research. Nucleic Acids Res. 44, e108–e108 (2016).

**Benchmark curation decision**:

- Finally, using the KEEP/?/REMOVE column from the *mosaic benchmark curation sheet*, we declared:
    - If our procedure defined a variant as TP and it was marked as KEEP, our verdict was to KEEP it.
    - If our procedure defined a variant as FP and it was marked as REMOVE, our verdict was to REMOVE it.
    - Otherwise, we perform a manual curation with IGV to complement our verdict in the cases of discordance between our verdict and the GIAB call.

**Team Name**: Boutros Lab, UCLA

**Team Members:** Mohammed Faizal Eeman Mootor, Yash Patel, Takafumi N. Yamaguchi, Paul C. Boutros

##### General Description of Methods

**Data Validation:**

All pipelines implemented in this project utilize PipeVal (v4.0.0-rc.2) to validate input and output files (Patel *et al*. 2024, PMID: 38341658).

**Alignment:**

Sequence reads were aligned to the GRCh38 reference genome, including decoy contigs (Broad Institute, 2016-07-21), using BWA-MEM2 (v2.2.1) (Li *et al.* 2019). The alignment process was conducted with default settings (i.e. without alternate-contig awareness). Duplicate reads were marked using Picard's MarkDuplicates (v3.0.0) (Picard Toolkit 2019). The Genome Analysis Toolkit (GATK) was used to perform Indel Realignment (v3.7.0) and Base Quality Score Recalibration (BQSR) (v4.2.4.1) (McKenna *et al.* 2010).

**Creating HG002-N (Normal) BAM:**

After BQSR, the AJ parental BAM files (HG003 and HG004) were merged using Picard's MergeSamFiles (v3.0.0). The merged BAM underwent header modification using SAMtools reheader (v1.15.1) to replace parental sample IDs (HG003, HG004) with an ID derived from the AJ son and designated as “HG002-N" (Danecek *et al.* 2021).

**Somatic Variant Calling:**

The BQSR BAM of the AJ son HG002 was treated as the tumor sample while the merged parental BAM, HG002-N was treated as the normal sample. Somatic variant calling was performed using the call-sSNV (v7.0.0) pipeline with the tumor/normal BQSR BAMs (Patel *et al*. 2024, PMID: 38341660). The pipeline has two main steps: (1) calling somatic variants using four different somatic variant callers: Mutect2 (v4.4.0.0), SomaticSniper (v1.0.5.0), Strelka2 (v2.9.10) and MuSE (v2.0.4) (McKenna *et al.* 2010, Larson *et al.* 2012, Kim *et al.* 2018, Fan *et al.* 2016), and (2) intersecting the resulting variant calls using BCFtools (v1.17) (Danecek *et al.* 2021) to produce consensus variants detected by at least two or more callers. Both consensus and individual caller variants were considered for the HG002 somatic mosaic benchmark evaluation. All alignment and varying calling steps were implemented in Nextflow-based pipelines (Patel *et al.* in preparation).

**Input data**:
For this benchmark evaluation, we used the GIAB AJ trio (HG002/HG003/HG004) Illumina PCR-free WGS (2x150bp, 300X) FASTQs.

**Read alignment and mapping**:

- Each replicate from each run was processed using the following steps:
    - Alignments (BWA-MEM2 v2.2.1 & SAMtools v1.12)

```
bwa-mem2 mem -R
\"@RG\\tID:${read_group_id}.Seq${lane}\\tCN:${sequencing_center}\\tLB:${library_id}\\tPL: ${platform_technology}\\tPU:${platform_unit}\\tSM:${sample}"
reference-GRCh38.fa R1.fastq R2.fastq | samtools view -S -b > lane.bam
```

    - Sort alignments from each lane BAM (SAMtools v1.15.1)

```
samtools sort -O bam -o sorted-lane.bam lane.bam
```

    - Merge lane BAMs for each sample (SAMtools v1.15.1)

```
samtools merge --write-index -o sample.bai sorted-lane{1..n}.bam
```

    - Mark duplicates (Picard v3.0.0)

```
java -jar picard.jar MarkDuplicates --VALIDATION_STRINGENCY LENIENT –INPUT
sample.bam –OUTPUT sample_markdup.bam --METRICS_FILE markdup_bam.metrics
--ASSUME_SORT_ORDER coordinate --PROGRAM_RECORD_ID MarkDuplicates --CREATE_INDEX
true
```

    - Indel realignment (GATK v3.7.0)

```
java -jar GenomeAnalysisTK.jar --analysis_type RealignerTargetCreator
sample_markdup.bam --reference_sequence Homo_sapiens_assembly38.fasta
```

```
--knownAlleles Mills_and_1000G_gold_standard.indels.hg38.vcf.gz --knownAlleles
Homo_sapiens_assembly38.known_indels.vcf.gz
--allow_potentially_misencoded_quality_scores --targetIntervals
sample_chr{n}.intervals --out sample_indelrealigned-chr{n}.bam --intervals
chr{n}-contig.interval_list
```

- Base quality score recalibration (GATK v4.2.4.1)

```
gatk BaseRecalibrator sample-indel-realign{1..n}.bam —-reference
reference-GRCh38.fa --verbosity INFO --known-sites
Mills_and_1000G_gold_standard.indels.hg38.vcf.gz --known-sites
Homo_sapiens_assembly38.known_indels.vcf.gz --known-sites
bundle_v0_dbsnp138.vcf.gz --output sample_recalibration_table.grp --read-filter
SampleReadFilter —-sample sample-id

gatk ApplyBQSR –input sample-indel-realign{n}.bam --bqsr-recal-file
sample_recalibration_table.grp –reference reference-GRCh38.fa --read-filter
SampleReadFilter —-output stdout --sample sample 2> .command.err | samtools view
-h | awk '(/^@RG/ && /SM:sample/) || ! /^@RG/' | samtools view -b -o
sample-bqsr-chr{n}.bam
```

- **Tools:**
  - BWA-MEM2 v2.2.1 & SAMtools v1.12
  - SAMtools v1.15.1
  - Picard v3.0.0
  - GATK v3.7.0 for Indel Realignment
  - GATK v4.2.4.1

**Normal Sample Creation (HG002-N):**

- **Merge HG003 and HG004 BAMs (Picard v3.0.0)**

```
java -jar picard.jar MergeSamFiles I=HG003.bam I=HG004.bam
O=HG003-HG004-merged.bam CREATE_INDEX=true
```

- **Reheader merged HG003-HG004 BAM to HG002-N (SAMtools v1.15.1)**

```
samtools reheader HG002-N.header HG003-HG004-merged.bam > HG002-N.bam
```

- **Tools**
  - Picard v3.0.0
  - SAMtools v1.15.1

**Mosaic variant calling**:

- **Variant calling tool(s) – mode: From the call-sSNV v7.0.0 pipeline**
  - MuSE v2.0.4

```
MuSE call -f reference-GRCh38.fa -O MuSE-HG002-T HG002-T.bam HG002-N.bam
MuSE sump -I MuSE-HG002-T.txt -G -O MuSE-HG002-T-raw.vcf -D dbsnp.vcf.gz
```

- Mutect2 v4.4.0.0

[GATK Mutect2 Workflow]

- SomaticSniper v1.0.5.0 (downstream filtering not included below)

```
bam-somaticsniper -q 1 `# map_qual 1 is recommended` -Q 15 `# somatic_qual
default to 15` -T 0.85 `# theta default to 0.85` -N 2 `# haplotypes default to
2` -r 0.001 `# prior_haplotypes default to 0.001` -F vcf `# output_format here
is vcf` `# The next 2 lines are included because in the original script
'use_prior_prob' was turned on` -J -s 0.01 -f reference-GRCh38.fa HG002-T.bam
HG002-N.bam SomaticSniper-HG002-T.vcf
```

- Strelka2 v2.9.10 + Manta v1.6.0

```
configureStrelkaSomaticWorkflow.py –normalBam HG002-N.bam –tumorBam HG002-T.bam
--referenceFasta reference-GRCh38.fa –indelCandidates Manta-Indel-candidates.vcf
--runDir StrelkaSomaticWorkflow
```

- BCFtools v1.17 to create consensus somatic calls

```
bcftools isec --nfiles +2 --output-type z --prefix isec-2-or-more ${vcf-list}
bcftools --output-type v --output BCFtools-HG002-T_SNV-concat.vcf
--allow-overlaps --rm-dups all ${vcf-list-from-above-step}
```

- **Tools:**
  - MuSE v2.0.4
  - GATK v4.4.0.0 (Mutect2)
  - SomaticSniper v1.0.5.0
  - Strelka2 v2.9.10 and Manta v1.6.0
  - BCFtools v1.17

**Benchmark curation decision**:

- Finally, using the KEEP/?/REMOVE column from the *mosaic benchmark curation sheet*, we declared:
  - If our procedure defined a variant as TP and it was marked as KEEP, our verdict was to KEEP it.
  - If our procedure defined a variant as FP and it was marked as REMOVE, our verdict was to REMOVE it.
  - Otherwise, we perform a manual curation with IGV to complement our verdict in the cases of discordance between our verdict and the GIAB call.
